# B3GALT6 Promotes Dormant Breast Cancer Cell Survival and Recurrence by Enabling Heparan Sulfate-Mediated FGF Signaling

**DOI:** 10.1101/2023.01.31.526529

**Authors:** Amulya Sreekumar, Michelle Lu, Biswa Choudhury, Tien-chi Pan, Dhruv K. Pant, Christopher J. Sterner, George K. Belka, Takashi Toriumi, Brian Benz, Matias Escobar-Aguirre, Francesco E. Marino, Jeffrey D. Esko, Lewis A. Chodosh

## Abstract

Breast cancer mortality results primarily from incurable recurrent tumors seeded by dormant, therapy-refractory residual tumor cells (RTCs). Understanding the mechanisms enabling dormant RTC survival is therefore essential for improving patient outcomes. We derived a dormancy-associated RTC signature that mirrors the transcriptional response to neoadjuvant chemotherapy in patients and is enriched for extracellular matrix-related pathways. In vivo CRISPR-Cas9 screening of dormancy-associated candidate genes identified the galactosyltransferase B3GALT6 as a functional regulator of RTC fitness. B3GALT6 covalently attaches glycosaminoglycans (GAGs) to proteins to generate proteoglycans and its germline loss-of-function causes skeletal dysplasias. We determined that B3GALT6-mediated biosynthesis of the GAG heparan sulfate predicts poor patient outcomes, promotes tumor recurrence by enhancing dormant RTC survival in multiple contexts, and does so via a B3GALT6-heparan sulfate/HS6ST1-heparan 6-*O*-sulfation/FGF1-FGFR2 signaling axis. These findings identify a role for B3GALT6 in cancer and suggest targeting FGF signaling as a novel approach to preventing recurrence by eradicating dormant RTCs.

## Introduction

Despite advances in early detection and treatment, breast cancer remains the leading cause of cancer-related deaths among women worldwide^1^. Mortality results predominantly from incurable recurrences that arise years, or even decades, following treatment of the primary tumor (PT)^2,3^. Since recurrent tumors derive from residual tumor cells (RTCs) that survive therapy, and are believed to reside in a reversibly quiescent, non-proliferative state of cellular dormancy^4^, depleting this critical pool of cells by targeting their survival mechanisms represents an attractive approach to preventing breast cancer recurrence.

RTC dormancy can be selected for or induced by exposure to therapy and/or interaction with a foreign microenvironment following cancer cell dissemination from the primary site. Experimental models recapitulating these paradigms of dormancy induction have been used to identify mechanisms underlying RTC persistence and recurrence. These include i) therapy-associated models that mimic RTC survival following targeted therapy by downregulating oncogenes such as *Her2*^5,6^, *Wnt1*^6,7^, and *Fgfr1*^8^, and ii) models that recapitulate interactions between RTCs and a foreign microenvironment, such as the D2.OR/D2A1 paired cell line model^9–11^.

Recent data suggest marked overlap of dormant RTC gene expression profiles that may be independent of both the location of RTCs at primary or metastatic sites and the stimulus responsible for dormancy entry^6,12^. Consequently, investigating conserved elements of dormancy-associated gene expression signatures may provide a tractable approach for identifying unique dependencies of dormant RTCs that could be targeted to induce their elimination, thereby preventing recurrence. Among these dependencies, crosstalk between RTCs and components of the extracellular matrix (ECM), including thrombospondin^13^, laminins^14,15^, fibronectin^16–19^, and collagens^20–22^, have been reported to promote RTC survival and dormancy. In contrast, little is known about the role of other ECM components, including proteoglycans, in regulating dormancy and RTC fitness.

Proteoglycans consist of unique core proteins covalently linked to one or more glycosaminoglycan (GAG) side chains^23^, and are subclassified as heparan sulfate, chondroitin/dermatan sulfate, or keratan sulfate proteoglycans based on the type of GAG they contain^24^. Following their assembly, proteoglycans can be embedded into the plasma membrane, secreted from the cell, or shed into the ECM where they orchestrate a variety of biological activities^25,26^.

A defining feature of GAGs that underlies their biological function is their highly anionic nature^23^, which is imparted by their component uronic acid residues and by dynamic, spatially regulated sulfation^27^. Anionic GAGs avidly bind growth factors, thereby regulating cellular signaling as well as phenotypes such as proliferation and survival^23^. The impact of these modified glycans on RTC fitness and breast cancer progression is unknown.

Here, we performed an in vivo CRISPR-Cas9 screen targeting a core set of dormancy-associated, tumor cell-autonomous genes to identify functional regulators of dormant RTC survival. The strongest hit from this screen was *B3galt6*, a galactosyltransferase that is essential for proteoglycan assembly^28^ and whose germline loss-of-function gives rise to the connective tissue disorders spondyloepimetaphyseal dysplasia with joint laxity, type 1 (SEMD-JL1), spondylodysplastic Ehlers-Danlos syndrome (spEDS), and Al-Gazali syndrome^29–31^. A role for B3GALT6 in cancer has not been reported. Our studies identify proteoglycan synthesis as a critical vulnerability of dormant RTC fitness that promotes their survival and recurrence, in part through B3GALT6-enabled heparan sulfate proteoglycan synthesis that potentiates FGF signaling in RTCs. Our observations indicate that RTC survival is dependent upon tumor cell-autonomous heparan sulfate synthesis and FGF signaling, which may constitute a targetable node to prevent lethal breast cancer recurrences.

## Results

### Dormant tumor cells display cell-autonomous upregulation of ECM-related transcripts following therapy

To understand the mechanisms underlying dormant RTC survival and persistence, we have developed and characterized doxycycline-inducible genetically engineered mouse (GEM) models of breast cancer^5–7,32–37^. These models of cellular dormancy^6^ enable robust spatiotemporal regulation of oncogene activation, thereby modeling the effects of targeted therapy in patients and permitting study of the impact of oncogenic pathway inhibition in genetically complex PTs, as well as the contribution of dormant RTCs to spontaneous recurrence.

RTCs that survive PT regression induced by Her2 downregulation in *MMTV-rtTA;TetO-Her2/neu* (*MTB/TAN*) mice^5^, or by Wnt1 downregulation in *MMTV-rtTA;TetO-Wnt1* (*MTB/TWNT*) mice^7^, were previously isolated to generate RTC-specific gene expression signatures^6^.

Dormancy signatures from *MTB/TAN* and *MTB/TWNT* mice were highly concordant, suggesting conserved elements of the dormant state^6^. Dormancy-associated changes in gene expression in RTCs in vivo in the context of therapy could be imposed by the microenvironment or could result from cell-autonomous determinants. To identify a core set of tumor cell-autonomous regulators of RTC survival and dormancy, we cultured PT cells (PTCs) from the Her2-dependent model in vitro (**Fig. 1A**). Maintaining Her2-dependent tumor cells in the presence of doxycycline under reduced serum conditions resulted in high levels of Her2 expression and proliferation as measured by Ki67 and EdU (D0; *baseline*) (**Fig. 1B, C**). Withdrawing doxycycline elicited the rapid decay of Her2 levels (Her2 *deinduction*) and cellular proliferation, with negligible levels of Ki67 and EdU detected at 7, 14, and 28 days post-doxycycline withdrawal (D7, D14, D28). Importantly, re-addition of doxycycline to quiescent tumor cells (Her2 *reinduction*) at each of these time points rapidly restored Her2 levels and proliferation to baseline levels (D7+, D14+, D28+) (**Fig. 1B, C**). Thus, the quiescent state of these cells is reversible, which is a *sine qua non* of cellular dormancy, confirming that this in vitro system recapitulates key features of dormancy observed in vivo^6,32,38^.

**Fig. 1:**
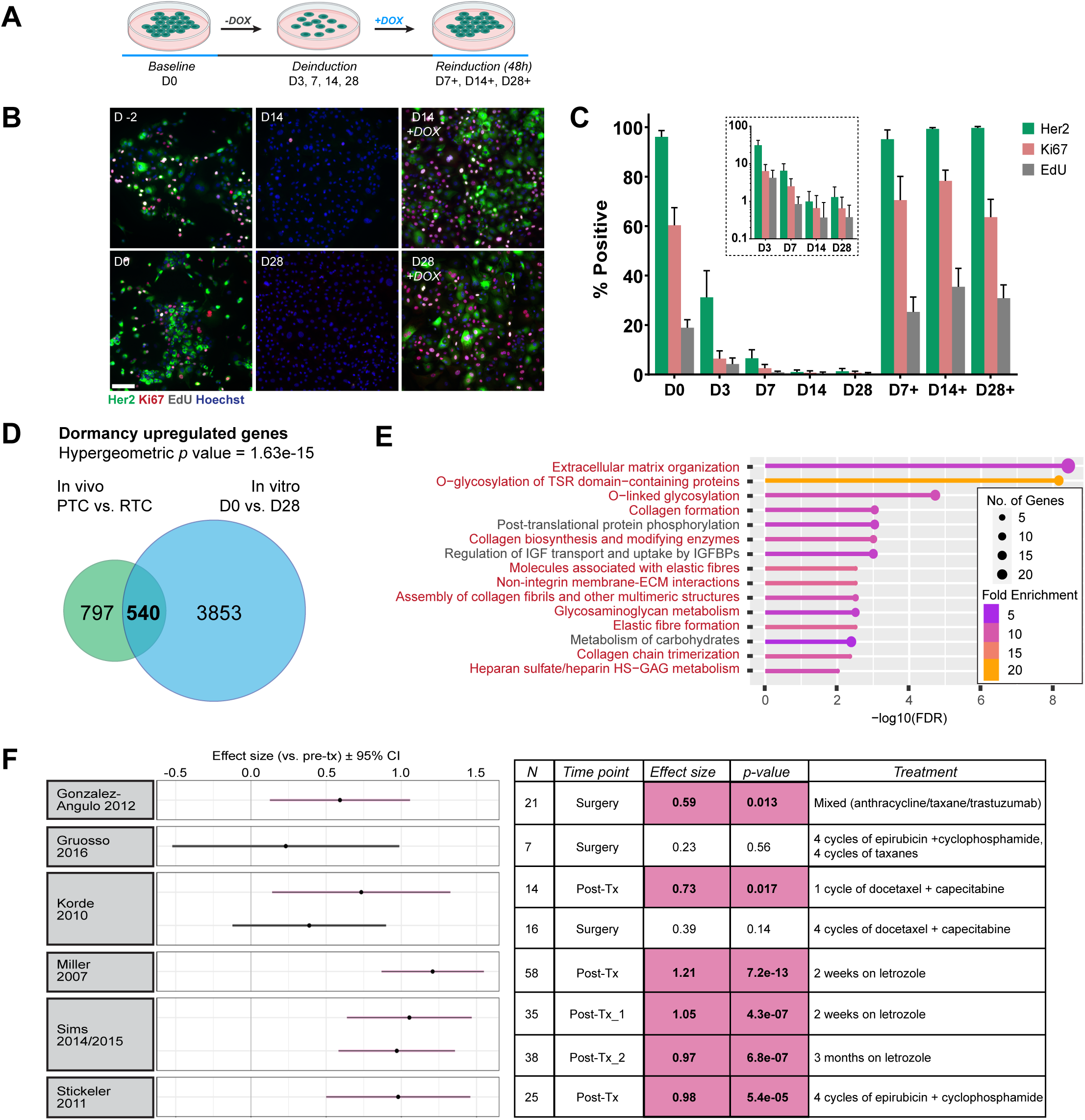
Dormant tumor cells display cell-autonomous upregulation of ECM-related transcripts following therapy. **A.** Experimental schematic indicating time points for in vitro dormancy (IVD) assay using *MTB/TAN* cells. **B.** Immunofluorescence for Her2 (green), Ki67 (red), and EdU (grey) with Hoechst nuclear staining (blue) during IVD. **C.** Quantification of (**B**) represented as mean ± standard deviation (SD). Inset displays the percentages at deinduction time points on a log_10_ y-axis. Scale bar=100µm. **D.** Hypergeometric test and Venn diagram depicting the overlap between the in vivo and IVD-derived upregulated gene set. **E.** Top 15 Reactome gene ontology terms for the overlapping upregulated gene set. ECM-associated categories highlighted in red. **F.** Core RTC signature enrichment following neoadjuvant therapies vs. pre-treatment (pre-tx) samples across 6 patient datasets measured by effect size (mean of pair-wise difference in signature scores). Significant effects are highlighted in pink.

To identify a core set of dormancy-associated genes, we performed RNA-sequencing on cells in vitro at *baseline*, *deinduction*, and *reinduction* time points. We evaluated whether dormancy-associated gene expression changes are conserved in dormant *MTB/TAN* and *MTB/TWNT* RTCs in vivo (PTCs vs. D28 RTCs) and *MTB/TAN* cells in vitro (D0 vs. D28) and found extensive overlap in the set of downregulated genes between these models (*p*=6.87e-21) (**Fig. S1A**). As anticipated, gene ontology analysis of the overlapping set of 747 downregulated genes identified enrichment for pathways related to cell cycle, cellular biosynthesis, and translation (**Fig. S1B**).

Conversely, we found a highly significant overlap among genes upregulated during dormancy in vivo (PTCs vs. D28 RTCs) and in vitro (D0 vs. D28) (540 genes, *p*=1.63e-15) (**Fig. 1D**). Intriguingly, gene ontology analysis of this overlapping gene set identified enrichment for multiple pathways related to the organization and metabolism of ECM components (**Fig. 1E**). We refer to the overlapping in vivo and in vitro-derived up- and down-regulated gene sets as the core RTC signature.

Thus, by integrating dormancy-associated gene expression changes that occur in vitro and in vivo, we generated a therapy-associated core RTC signature that is enriched for conserved features of the dormant state, including the tumor cell-autonomous synthesis of ECM components.

### A core RTC dormancy signature recapitulates neoadjuvant chemotherapy-associated gene expression changes and predicts favorable outcomes in patients

To evaluate whether the core RTC signature exhibited by dormant tumor cells following Her2 downregulation is clinically relevant, we interrogated data from six publicly available datasets comprised of paired gene expression profiles for primary breast cancers and residual tumors in patients that were first biopsied and then surgically resected following neoadjuvant chemotherapy^39–45^. Five of these six datasets displayed significant enrichment for the core RTC signature post-therapy (**Fig. 1F**), even after excluding proliferation-associated genes (**Fig. S1C**). Notably, enrichment for this signature was independent of both the type (chemotherapy or endocrine therapy) and duration of therapy. These findings indicate that mouse dormancy models recapitulate clinically-relevant aspects of the response of breast cancers to therapy in patients and suggest that therapy-refractory tumor cells that persist in patients exhibit changes in gene expression that are similar to those associated with cellular dormancy in mice.

To evaluate the prognostic power of the core RTC signature in breast cancer patients, we performed a meta-analysis of patient-derived recurrence data. Analogous to prior observations for an in vivo *MTB/TAN* and *MTB/TWNT*-derived RTC dormancy signature^6^, we found that patients whose PTs were enriched for the core RTC signature exhibited a striking decrease in risk of recurrence (HR=0.14; *p*=1.1e-29), ostensibly reflecting a higher propensity of such tumors to display indolent properties (**Fig. S1D**). This association was substantially stronger for the core RTC signature (HR=0.14) vs. the in vivo signature alone (HR=0.49)^6^, and persisted following the removal of proliferation-associated genes (*p*=3.2e-18), indicating that the core signature reflects features of cellular dormancy beyond those related to cell cycle-arrest (**Fig. S1E**). Furthermore, since >75% of patient tumors in this dataset were estrogen receptor positive (ER+) and displayed metastatic recurrence, our findings suggest that a conserved RTC signature generated from ER-negative RTCs at local sites can inform the study of mechanisms enabling RTC survival and persistence in the contexts of both distant recurrence and ER+ disease^6^.

In conclusion, we found enrichment of a core RTC signature in residual lesions from patients treated with different neoadjuvant chemotherapies, and determined that this signature is significantly associated with risk of tumor recurrence in patients across multiple breast cancer subtypes, and at both local and distant sites of recurrence.

### An in vivo ECM-focused loss-of-function screen identifies *B3galt6* as a regulator of RTC fitness

Having derived a core RTC signature, we reasoned that genes upregulated during dormancy in a manner that is reversible following Her2 reinduction might be involved in promoting RTC dormancy and/or survival. To test this hypothesis, we focused on a clustered set of genes enriched for ECM-associated gene ontology terms that displayed a similar temporal pattern of upregulation during in vitro dormancy (IVD) that was reversed upon Her2 reinduction (**Fig. 2A**).

**Fig. 2:**
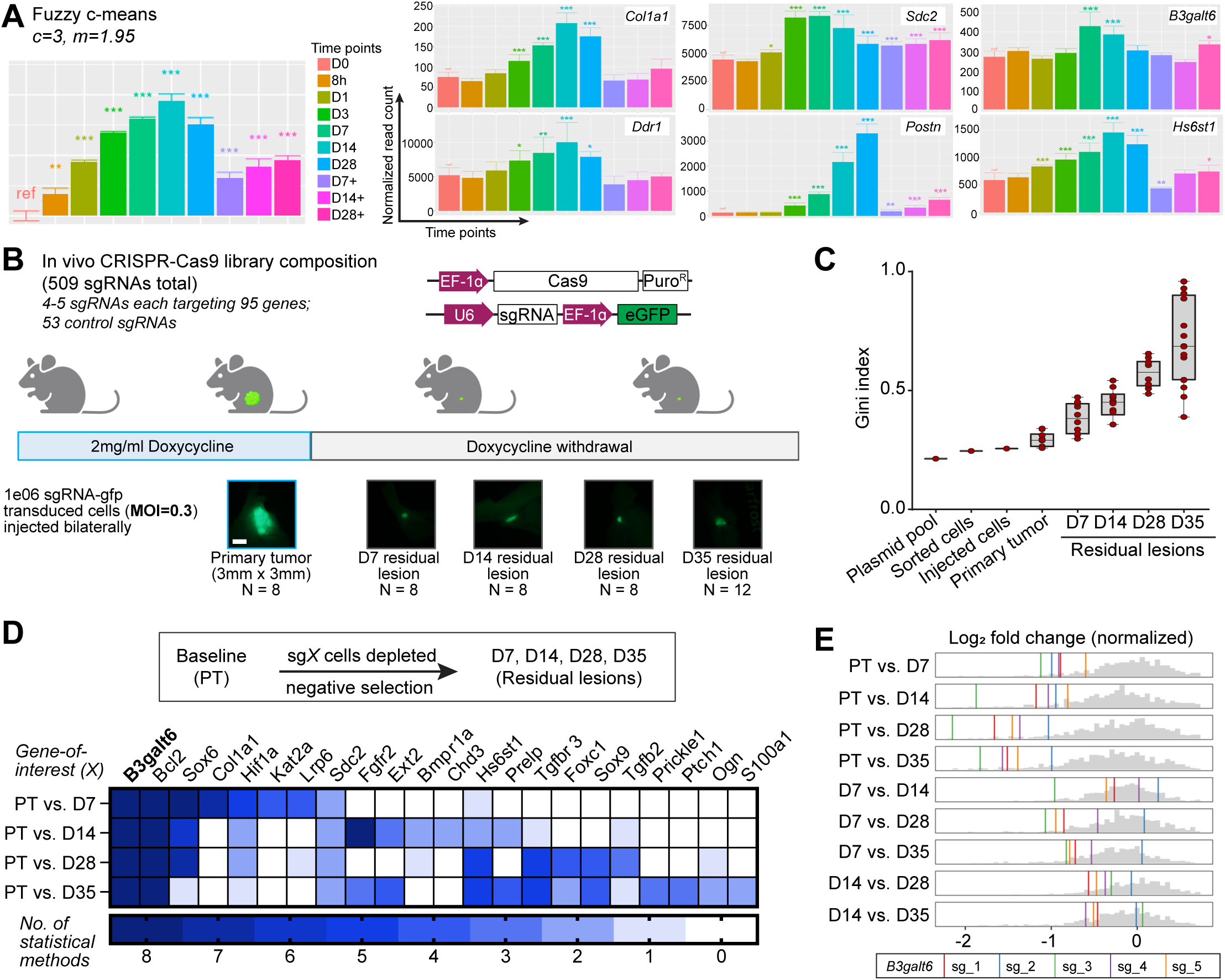
An ECM-focused loss-of-function screen identifies *B3galt6* as a novel regulator of RTC fitness in vivo. **A.** Fuzzy c-means clustering using cluster # (c)=3, fuzzifier (m)=1.95 identifies one cluster that displays reversible, dormancy-dependent upregulation of genes at 8h, D1, D3, D7, D14, D28 (deinduction) time points. 6 genes (of 3450) in this cluster are indicated. Asterisks indicate significant changes in normalized read counts vs. D0 (baseline) \**p*<0.05, \*\**p*<0.01, \*\*\**p*<0.001, \*\*\*\**p*<0.001. **B.** In vivo CRISPR-Cas9 screen schematic. Stereoscope images of representative lesions are shown. Scale bar=2mm. **C.** Gini index for heterogeneity at sequential time points assayed in the screen. Data are represented as median ± range. **D.** Target identification criterion and list of CRISPR-Cas9 screen depletion hits identified by 2-8 statistical methods used for calling hits. **E.** Log_2_ fold change (normalized) gene-level effect size plots for *B3galt6* sgRNAs in different pairwise comparisons using PT, D7, or D14 as baselines. Grey histogram depicts the background distribution of sgRNAs in the CRISPR-Cas9 screen, colored vertical lines depict each sgRNA targeting *B3galt6*.

To determine whether genes within this cluster might functionally regulate RTC survival in a tumor cell-intrinsic manner, we performed an in vivo CRISPR-Cas9-based loss-of-function screen. We designed a custom sgRNA library with 4-5 sgRNAs targeting each of 95 candidate genes that displayed strong, dormancy-selective expression and were enriched within ECM-associated ontology terms (**Table S1**). An additional 53 sgRNAs comprising positive controls anticipated to be lethal to proliferative cells (*e.g*., sg*Rpa3*, sg*Pcna*) and negative controls consisting of non-targeting sgRNAs and sgRNAs targeting inert sites (*e.g.*, sg*Rosa*) anticipated to lack selection during tumor dormancy and recurrence (**Fig. 2B, Table S1**).

*MTB/TAN-*derived Her2-dependent PTCs constitutively expressing Cas9 (Her2-dependent-Cas9 cells) were transduced with this GFP-labeled sgRNA library at low multiplicity of infection (MOI=0.3) and sorted to ensure that each cell contained a single sgRNA. After confirming that all sgRNAs were represented within GFP+ sorted cells (**Fig. S2A**), cells were orthotopically injected into nude (*nu/nu*) mice. To assess sgRNA selection during disease progression, we allowed Her2-driven PTs to form in the presence of doxycycline, then withdrew doxycycline to induce tumor regression to a non-palpable, dormant state (**Fig. S2B)**. PTs and residual lesions (RLs) were harvested at time points corresponding to early (D7), mid (D14) and late (D28, D35) dormancy following tumor regression to assess sgRNA composition at each stage of tumor progression (**Fig. 2B**).

By sequencing the initial plasmid pool, sorted cells, injected cells, PT and RL samples, we confirmed that all sgRNAs were detectable at each time point (**Fig. S2A**). Calculating the Gini index to quantify the skewness of sgRNA distribution revealed that the plasmid pool, sorted cells, and injected cells displayed low Gini indices (median=0.21, 0.25, and 0.26, respectively), confirming that the distribution of sgRNAs remained relatively homogeneous at these points. PTs displayed a slightly higher Gini index than pre-injection samples (median=0.29), suggesting only modest selection of sgRNAs in the presence of the strong oncogenic driver, Her2. In contrast, RLs displayed substantial stepwise increases in Gini indices at D7, D14, D28, and D35 (median=0.38, 0.45, 0.58, 0.69, respectively), confirming that sgRNAs were progressively selected throughout the dormancy period (**Fig. 2C**).

To identify sgRNAs that underwent selection, we employed 8 analytical methods and ranked sgRNAs identified by 2 or more methods. Consistent with our prior findings that RTCs in this model are non-proliferative following Her2 downregulation^6^, sg*Rpa3* and sg*Pcna* positive controls were depleted during PT formation (**Fig. S2C**), but not following Her2 downregulation (**Fig. S2D**).

We reasoned that sgRNAs targeting genes that maintain cell cycle-arrest following Her2 down-regulation would be enriched in RLs vs. PTs. Our finding that *Ddr1,* a known regulator of cell cycle arrest^22^, is enriched in this manner lends credence to this hypothesis (**Fig. S2E**). Conversely, we anticipated that sgRNAs targeting genes that promote cell survival following Her2 down-regulation would be depleted in RLs vs. PTs, as supported by the identification of the pro-survival gene, *Bcl2*, as one of two top hits in this analysis (**Fig. 2D**).

The second top putative pro-survival hit identified by all 8 methods in each of four pairwise comparisons of PT and RL time points was *B3galt6* (**Fig. 2D, S2F**). Analyzing the distribution of sgRNAs targeting *B3galt6* (colored bars) vs. all sgRNAs (grey histogram) in each pairwise comparison confirmed that *B3galt6* sgRNAs were strongly depleted within 7 days following Her2 downregulation (PT vs. D7, D14) (**Fig. 2E, S2F**). Notably, *B3galt6* sgRNAs exhibited further depletion at later dormancy time points, as revealed by comparisons of early (D7) to later (D14, D28, D35) RL time points (**Fig. 2E, S2F**). This strong, persistent negative selection against cells harboring *B3galt6* sgRNAs suggests a potential role for B3GALT6 in dormancy.

### B3GALT6 promotes RTC survival following therapy-associated dormancy

Beta-1,3-galactosyltransferase (B3GALT6) catalyzes the addition of the second Gal residue in the GlcA-<u>Gal</u>-Gal-Xyl-*O-*tetrasaccharide linker that is essential for the attachment of sulfated GAGs to proteoglycan core proteins^28^. Accordingly, B3GALT6 enzymatic activity is required for proteoglycan assembly. Based on their linear chains of repeating disaccharides, GAGs can be classified as: (i) heparan sulfate (GlcNAc-GlcA or GlcNAc-iduronic acid (IdoA)); (ii) chondroitin sulfate (GalNAc-GlcA); or (iii) dermatan sulfate (resulting from the epimerization of at least one GalNAc-GlcA to GalNAc-IdoA) (**Fig. 3A**). Consequently, deletion of the *B3galt6* gene using CRISPR-Cas9 should ablate both heparan and chondroitin/dermatan sulfate GAGs on cell surfaces^46^.

**Fig. 3:**
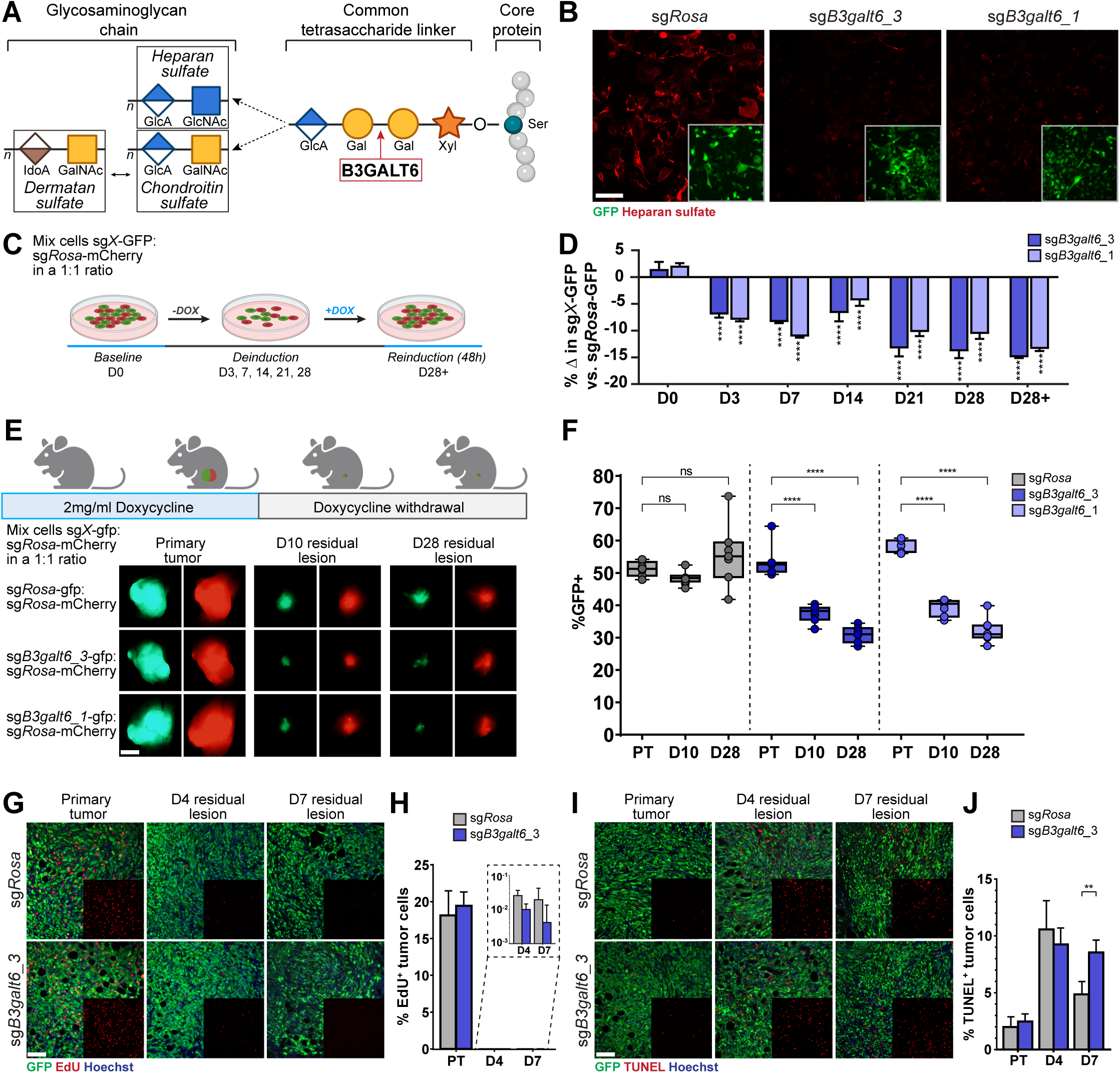
B3GALT6 promotes RTC survival during dormancy. **A.** B3GALT6 (red box) function in tetrasaccharide linker synthesis and proteoglycan assembly. Typical disaccharide repeat units in heparan sulfate, chondroitin sulfate, and dermatan sulfate are indicated. Ser=Serine, Xyl=Xylose, Gal=Galactose, GlcA=Glucuronic acid, GlcNAc=N-Acetylglucosamine, GalNAc=N-Acetylgalactosamine, IdoA=Iduronic acid. **B.** Immunofluorescence for heparan sulfate (red) in Her2-dependent-Cas9 cells transduced with sg*Rosa*, sg*B3galt6*_3, or sg*B3galt6*_1 sgRNAs (green). Scale bar=100µm. **C.** IVD competition assay schematic. **D.** ddPCR data quantifying percentage change in sg*B3galt6_3* (dark blue) and sg*B3galt6*_1 (light blue) GFP+ cells normalized to sg*Rosa* (grey) cell numbers. Data are represented as mean ± SD. \*\*\*\**p*<0.001. **E.** Schematic for in vivo competition assay with stereoscope images of representative lesions harvested. Scale bar=2mm. **F.** ddPCR data quantifying GFP+ cell percentage in sg*Rosa* (grey), sg*B3galt6*_3 (dark blue), and sg*B3galt6*_1 (light blue) groups. Data are represented as median ± range. ns=non-significant, \*\*\*\**p*<0.001. **G-J.** Immunofluorescence (**G**) and quantification (**H**) for EdU (red) or for TUNEL (red) (**I, J**) in Her2-dependent-Cas9 sg*Rosa* and sg*B3galt6*_3 (green) PTs, D4, and D7 residual lesions (RLs). Scale bar=100µm. Quantification in the sg*Rosa* (grey) and sg*B3galt6*_3 (dark blue) groups is represented as mean ± SD. \*\**p*<0.01.

To test this, we transduced Her2*-*dependent-Cas9 cells with sg*Rosa* or sg*B3galt6* vectors expressing GFP, performed ICE analysis^47^, and confirmed that sg*B3galt6* efficiently induced indels predicted to result in loss-of-function mutations in >85% of tumor cells (**Fig. S3A, B**). Immunofluorescence performed on sg*Rosa* and sg*B3galt6* cells using a heparan sulfate antibody demonstrated a marked reduction in heparan sulfate levels in sg*B3galt6* cells vs. sg*Rosa* controls, providing further confirmation that these sgRNAs functionally reduce B3GALT6 activity (**Fig. 3B**).

To independently validate findings from our CRISPR-Cas9 screen, an in vitro competition assay was performed in which sg*Rosa*-GFP and sg*B3galt6*-GFP transduced Her2-dependent-Cas9 cells were plated at a 1:1 ratio with sg*Rosa*-mCherry cells on D-3. Genomic DNA was harvested from cells at D-2 or D0 (*baseline*) in the presence of doxycycline, at D3, D7, D14, D21, and D28 following doxycycline withdrawal (*deinduction*), and at 48 hr after doxycycline re-addition to D28 deinduction cells (D28+, *reinduction*). Changes in the percentage of sg*B3galt6-*GFP:sg*Rosa*-mCherry cells relative to sg*Rosa*-GFP:sg*Rosa*-mCherry controls were quantified by droplet digital PCR (ddPCR) (**Fig. 3C**). Tumor cells expressing sg*B3galt6* were significantly and progressively depleted (*p*<0.0001) throughout dormancy beginning as early as 3 days following Her2 downregulation (**Fig. 3D**). These data indicate that B3GALT6 loss impairs RTC fitness in a tumor cell-autonomous manner.

To confirm that this dormancy-selective depletion of sg*B3galt6* cells also occurs in vivo, as predicted by our CRISPR-Cas9 screen, we performed a competition assay analogous to that performed in vitro. sg*Rosa*-mCherry Her2-dependent*-*Cas9 cells admixed with an equal number of sg*Rosa*-GFP or sg*B3galt6*-GFP cells were injected into mice maintained on doxycycline, following which PT (*baseline*) as well as RL samples at D10 (*early*) and D28 (*late*) following doxycycline withdrawal were harvested (**Fig. 3E**).

Samples were imaged using a stereoscope prior to processing for genomic DNA isolation and ddPCR. While no differences in GFP intensity were visible in PT samples across groups, decreases in GFP intensity for each of the sg*B3galt6* guides vs. sg*Rosa* controls were clearly evident in D10 and D28 RL samples (**Fig. 3E**). Consistent with this, ddPCR analysis confirmed pronounced negative selection against sg*B3galt6* tumor cells (*p*<0.0001) at D10 and D28 RL time points, as well as ongoing depletion from D10 to D28 (**Fig. 3F**). In contrast, sg*Rosa*-GFP cells showed no selection across time points. These data provide further evidence that B3GALT6 is required for maintaining RTC fitness in vivo.

To determine the cellular mechanisms by which B3GALT6 regulates RTC fitness in vivo, sg*B3galt6_*3-GFP or sg*Rosa*-GFP cells were injected into mice maintained on doxycycline, followed by harvest of PTs as well as early (D4, D7) RLs following doxycycline withdrawal. Mice were injected with EdU 2 hr prior to sacrifice. As anticipated, a dramatic decrease in cell proliferation was observed in control lesions D4 and D7 following doxycycline withdrawal, as revealed by visualizing incorporation of the S-phase marker EdU (**Fig. 3G, H**) and immunofluorescence for the cell cycle marker Ki67 (**Fig. S3C, D**). No differences were observed in the percentage of EdU+ or Ki67+ RTCs between sg*B3galt6* and sg*Rosa*-derived PTs or D4 or D7 RLs (**Figs. 3G, H; S3C, D**). These findings confirm that RTCs in both sg*Rosa* and sg*B3galt6* groups enter a quiescent state following Her2 downregulation, and that the observed depletion of sg*B3galt6* RTCs vs. control cells following Her2 downregulation is unlikely to result from B3GALT6-dependent differences in proliferation.

Next, we asked if the decrease in sg*B3galt6* vs. *sgRosa* RTCs following Her2 downregulation is a consequence of differential cell survival by quantifying apoptotic cells by terminal deoxynucleotidyl transferase dUTP nick end labeling (TUNEL) (**Fig. 3I, J**), and by immunofluorescence for cleaved caspase 3, (cc3) (**Fig. S3E, F**). While no differences were evident in the percentage of TUNEL+ or cc3+ in PTs or D4 RLs between sg*B3galt6* and *sgRosa* samples, a significant increase in the percentage of TUNEL+ (*p*=0.0009) and cc3+ tumor cells (*p*=0.033) was apparent at D7 in sg*B3galt6* cells vs. sg*Rosa* RL samples. Together, these data indicate that the impaired fitness observed in sg*B3galt6* tumor cells following Her2 downregulation is attributable to higher levels of apoptosis in dormant sg*B3galt6* RTCs.

### B3GALT6 promotes tumor recurrence following therapy-associated dormancy

In light of our observation that B3GALT6 promotes RTC fitness, we wished to determine whether this would impact the kinetics of spontaneous tumor recurrence. Her2-dependent-Cas9 cells transduced with sg*B3galt6*-GFP or sg*Rosa*-GFP at a target MOI=5 (**Fig. S4A, B**) were orthotopically injected into mice maintained on doxycycline to drive PT formation (**Fig. 4A**). sg*B3galt6* and sg*Rosa*-GFP groups displayed no differences in PT formation (*p*=0.26) (**Fig. S4C**).

**Fig. 4:**
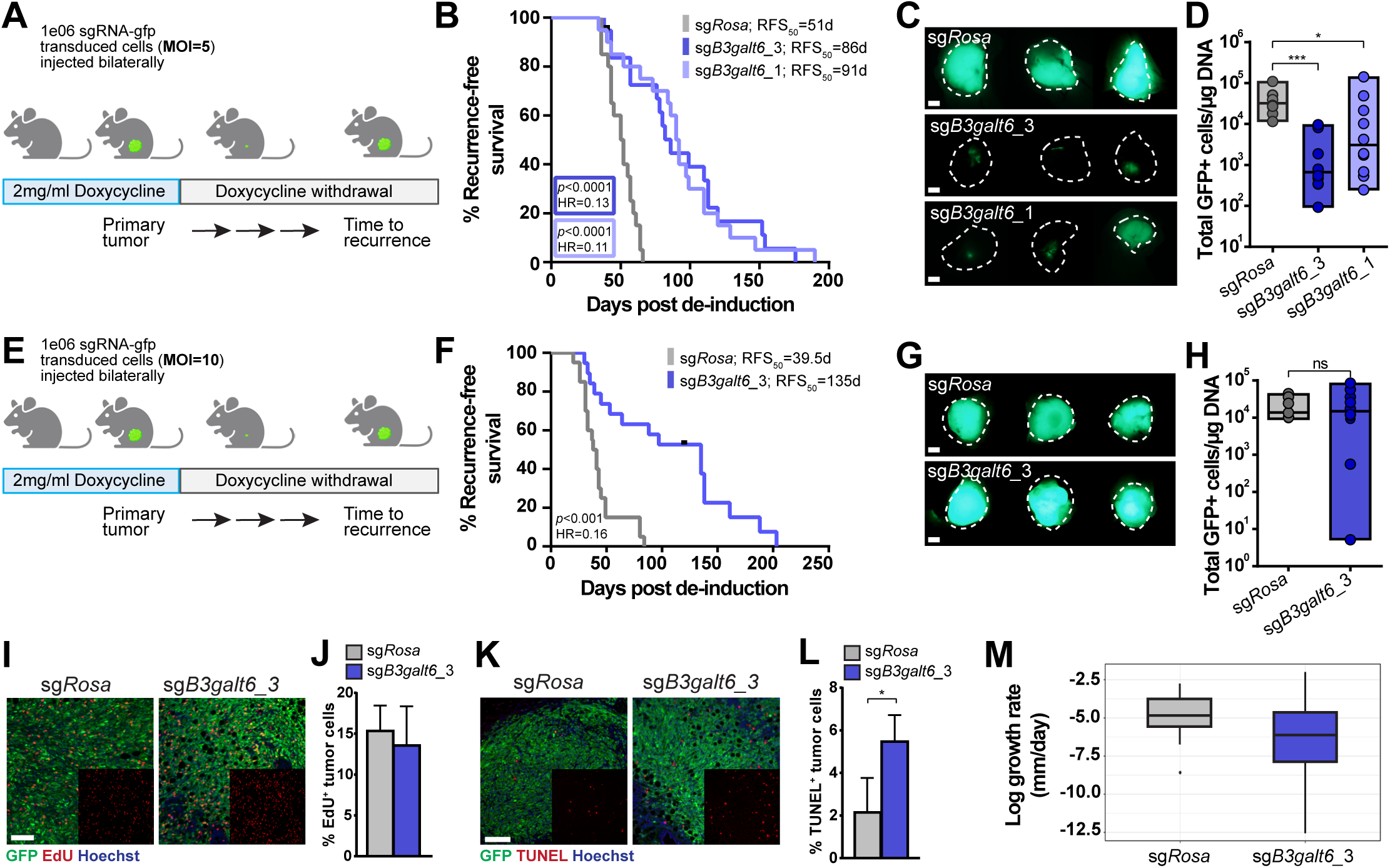
B3GALT6 promotes recurrence following dormancy. **A.** Recurrence-free survival assay using multiplicity of infection (MOI)=5. **B.** Kaplan-Meier analysis of recurrence-free survival for sg*Rosa* (grey), sg*B3galt6*_3 (dark blue), and sg*B3galt6*_1 (light blue) groups. RFS_50_=median time-to-recurrence. **C.** Stereoscope images of representative recurrences. Dotted white lines represent tumor edges identified in corresponding bright field images. Scale bar=2mm. **D.** Quantification of GFP+ cells in recurrent tumors as measured by ddPCR for sg*Rosa* (grey), sg*B3galt6*_3 (dark blue), and sg*B3galt6*_1 (light blue) groups. Data are represented as median ± range. \**p*<0.05, \*\**p*<0.001. **E.** Recurrence-free survival assay using MOI=10. **F.** Kaplan-Meier analysis of recurrence-free survival for sg*Rosa* (grey) and sg*B3galt6*_3 (dark blue) groups. RFS_50_=median time-to-recurrence. **G.** Stereoscope images of representative recurrences. Scale bar=2mm. **H.** Quantification of GFP+ cells measured by ddPCR for sg*Rosa* (grey) and sg*B3galt6*_3 (dark blue) recurrences. Data are represented as median ± range. ns=non-significant. **I-L.** Immunofluorescence (**I**) and quantification (**J**) for EdU (red) or for TUNEL (red) (**K, L**) in Her2-dependent-Cas9 sg*Rosa* and sg*B3galt6*_3 (green) recurrences. Scale bar=100µm. Quantification in sg*Rosa* (grey) and sg*B3galt6*_3 (dark blue) groups is represented as mean ± SD. \**p*<0.05. **M.** Log growth rate of sg*Rosa* (grey) and sg*B3galt6*_3 (dark blue) recurrent tumors. Data are represented as median ± interquartile range.

Following doxycycline withdrawal to induce tumor regression to a non-palpable, dormant state, mice were monitored for tumor recurrence. Strikingly, dormant RLs derived from sg*B3galt6*_3 (*p*<0.0001; HR=0.13) or sg*B3galt6*_1 (*p*<0.0001; HR=0.11) tumor cells displayed dramatically delayed median recurrence-free survival vs. sg*Rosa* controls (**Fig. 4B**). Moreover, whereas all 20 recurrent tumors in the sg*Rosa* control group were strongly GFP+, only 1/20 sg*B3galt6*_3 and 3/20 sg*B3galt6*_1 recurrent tumors were GFP+ (**Fig. 4C**, **S4D**). This strong negative selection against sg*B3galt6* GFP+ cells in recurrences was confirmed by ddPCR, which demonstrated >10-fold depletion of GFP+ cells in sg*B3galt6*_3 (*p*=0.0003) and sg*B3galt6*_1 (*p*=0.039) recurrences (**Fig. 4D**). Together, the marked delay in spontaneous tumor recurrence observed for sg*B3galt6*-cells, coupled with the observation that the small fraction (∼10%) of untransduced GFP-negative cells present efficiently and reproducibly outcompeted GFP+ sg*B3galt6* tumor cells during recurrent tumor formation, indicate that B3GALT6 is required for tumor recurrence.

To ensure that recurrences maintained B3GALT6 deletion (*i.e.*, remain GFP+), we transduced Her2-dependent-Cas9 cells with sg*Rosa*-GFP or sg*B3galt6*-GFP at a higher target MOI of 10 (**Fig. 4E**), which resulted in a transduction efficiency of >95% (**Fig. S4E, F**). Under these conditions, sg*B3galt6*-GFP cells exhibited a dramatic delay in median recurrence-free survival (135d) vs. sg*Rosa*-GFP (39.5d) cells (*p*<0.0001; HR=0.16) (**Fig. 4F**) that was even greater than that observed for cells transduced at lower MOI (86-91d vs. 51d) (**Fig. 4B**). Consistent with the higher MOI employed, nearly all recurrences arising from sg*B3galt6*-GFP (12/14) or sg*Rosa*-GFP (18/19) cells displayed strong GFP-positivity (**Fig. 4G, S4G**) and ddPCR confirmed that the number of GFP+ cells in sg*Rosa* and sg*B3galt6* recurrences was comparable (**Fig. 4H**). Further, immunofluorescence on recurrent tumor tissue confirmed that heparan sulfate levels were decreased in sg*B3galt6* vs. sg*Rosa* recurrences, as anticipated, thereby confirming that loss of B3GALT6 function was maintained in sg*B3galt6* recurrences (**Fig. S4H**). These observations suggest that transducing tumor cells with sg*B3galt6* at high MOI eliminated a bypass pathway in which untransduced (*i.e.*, GFP-negative) cells give rise to recurrence.

EdU labeling failed to identify differences in proliferation between sg*Rosa* and sg*B3galt6* recurrences (**Fig. 4I, J**). In contrast, TUNEL staining revealed an increase in apoptotic tumor cells in sg*B3galt6* vs. sg*Rosa* recurrent tumors (*p*=0.018) (**Fig. 4K, L**). Consistent with these findings, a trending, albeit non-significant, decreased growth rate was observed for sg*B3galt6* vs. sg*Rosa* recurrences (*p*=0.073) (**Fig. 4M**). Together, these findings indicate that B3GALT6 is required for efficient tumor recurrence from dormant residual disease post-therapy.

### B3GALT6 promotes tumor cell survival and outgrowth in microenvironment-induced dormancy

Our data to this point identified B3GALT6 as a critical regulator of RTC survival and recurrence in the context of therapy-associated dormancy. We next asked whether B3GALT6 might also play a functional role in microenvironment-induced dormancy. To address this question we used the D2.OR-D2A1 system of paired cell lines that share a common origin and grow comparably in two-dimensional (2D) culture, but manifest divergent growth properties (*i.e.*, D2.OR: dormant/indolent vs. D2A1: proliferative/aggressive) when cultured in 3D or at metastatic sites in vivo^16,20,48^.

We first determined whether a previously derived dormancy-associated gene expression signature generated by comparing D2.OR to D2A1 cells grown in 3D culture^48^ exhibited overlap with dormancy-associated gene expression changes in *MTB/TAN* cells following Her2 downregulation in vitro and in vivo. Gene expression changes observed in Her2-dependent tumor cells revealed strong and progressive enrichment of the D2.OR/D2A1 dormancy signature over the 28-day course following Her2 downregulation in vitro (**Fig. 5A**, *p*=1.88e-09). This enrichment was reversible following doxycycline readdition, which reactivates Her2 signaling and induces dormant tumor cells to re-enter the cell cycle (**Fig. 5A**). The D2.OR-derived dormancy signature was also enriched in a dormancy-specific manner in dormant *MTB/TAN* and *MTB/TWNT* RTCs in vivo 28 days following oncogene downregulation vs. either PTCs (*p*=7.4e-05, 1.7e-05, respectively) or recurrent tumor cells (*p*=6.2e-04, 1.1e-03, respectively) (**Fig. S5A**).

**Fig. 5:**
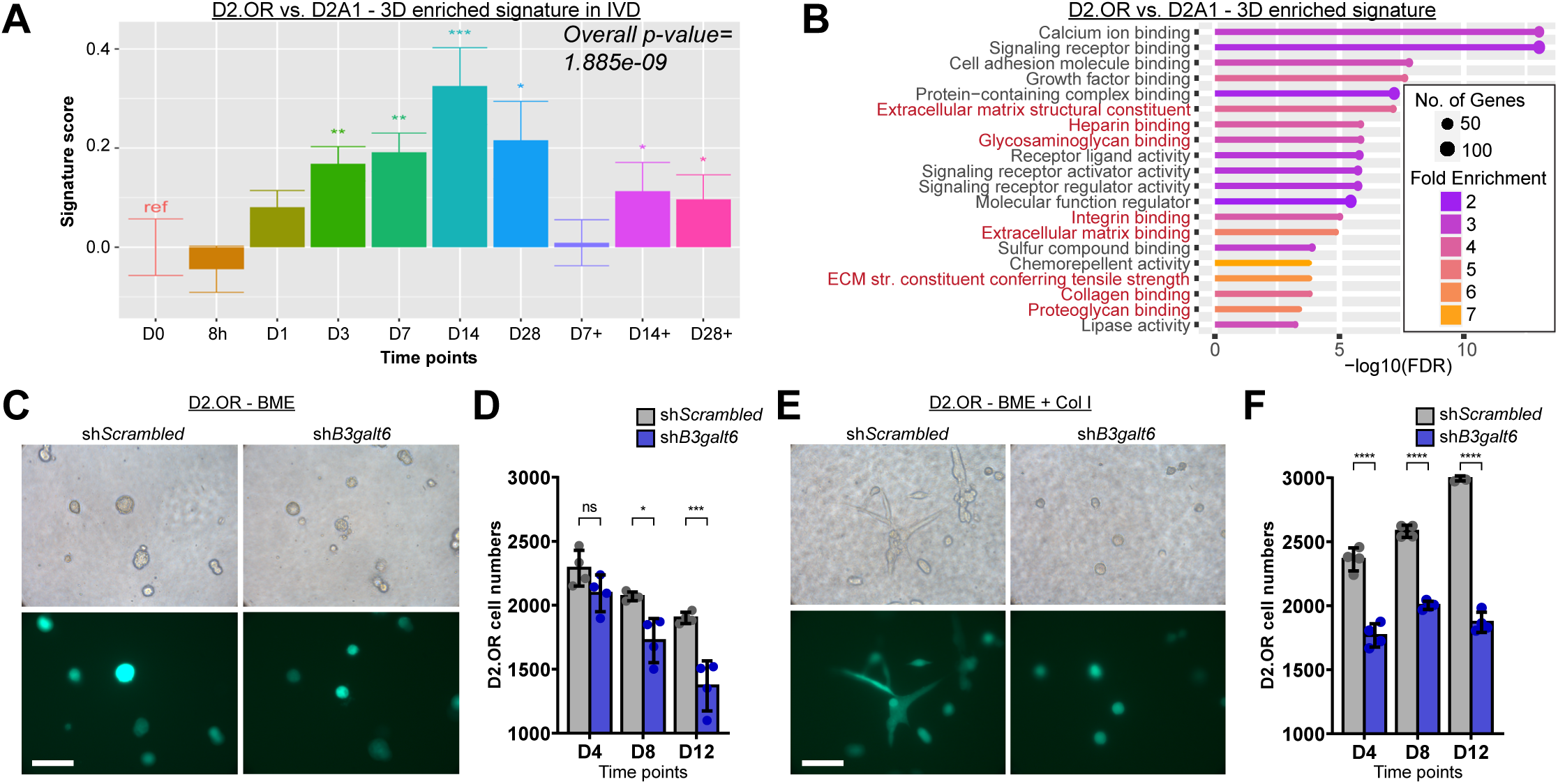
B3GALT6 promotes tumor cell survival and outgrowth in microenvironment-induced models of dormancy. **A.** Application of a gene expression signature derived from D2.OR (indolent) cells vs. D2A1 (aggressive) cells in 3D to IVD temporal profiling of Her2-dependent tumor cells. Asterisks indicate significant changes in normalized read counts vs. D0 (*baseline*) \**p*<0.05, \*\**p*<0.01, \*\*\**p*<0.001. **B.** Top 20 gene ontology terms for the overlapping upregulated set of 1018 genes. ECM-associated categories are highlighted in red. **C.** Brightfield and fluorescence images of sh*Scrambled* and sh*B3galt6* D2.OR cells grown in 3D on basement membrane extract (BME) and **D.** viable cell numbers measured at D4, D8, and D12 time points. Scale bar=100µm. Data are represented as mean ± SD. ns=non-significant, \**p*<0.05, \*\*\*\**p*<0.0001. **E.** Brightfield and fluorescence images of sh*Scrambled* and sh*B3galt6* D2.OR cells grown in 3D on BME + Col I and **F.** viable cell numbers measured at D4, D8, and D12 time points. Scale bar=100µm. Data are represented as mean ± SD. ns=non-significant, \**p*<0.05, \*\*\*\**p*<0.0001.

Next, we asked what gene ontology terms were enriched within the D2.OR-derived dormancy signature. As observed for the core RTC signature derived from therapy-associated dormancy models, the D2.OR dormancy signature was enriched for multiple pathways relating to ECM, proteoglycan, and GAG-related gene ontology terms (**Fig. 5B**, highlighted in red). These data indicate that dormancy-associated gene expression signatures are strongly enriched for ECM-related pathways in general, and proteoglycan and glycosaminoglycan-related pathways in particular, irrespective of whether the dormant state was induced by the microenvironment or associated with therapy.

D2.OR cells persist as dormant single cells over the course of 12 days when grown on basement membrane extract (BME), whereas D2A1 cells generate proliferative, spindle-shaped outgrowths under these same conditions^20^. Addition of Collagen I (Col I) to BME induces dormant D2.OR cells to resume proliferation and generate D2A1-like spindle-shaped colonies^20^. Using this system, we asked whether the dormant behavior of D2.OR cells in 3D culture is regulated by B3GALT6-mediated proteoglycan synthesis.

D2.OR and D2.A1 cells transduced with a sh*B3galt6* hairpin exhibited >80% knock-down of *B3galt6* transcripts (**Fig. S5B**). D2.OR cells transduced with sh*B3galt6* or control sh*Scrambled* hairpins were then overlaid on BME with or without addition of Col I. While the numbers of viable sh*B3galt6* and sh*Scrambled* cells each decreased over the 12-day course, *B3galt6* knockdown resulted in progressive and marked decreases in cell number at D8 (*p*=0.022) and D12 (*p*=0.0004) compared to control cells (**Fig. 5C, D**).

As anticipated, sh*Scrambled* D2.OR cells plated on BME + col I displayed increased viable cell numbers over this same time course (**Fig. 5E, F**). *B3galt6* knockdown in D2.OR cells yielded marked reductions in the numbers of viable tumor cells at D4, D8, and D12 compared to (*p*<0.0001) sh*Scrambled* controls. Indeed, the Col I-induced increase in number of D2.OR cells was entirely abrogated by *B3galt6* knockdown, as viable D2.OR sh*B3galt6* cells failed to increases in number when grown under these conditions (**Fig. 5E, F**). Additionally, while sh*Scrambled* D2.OR cells formed spindle-like colonies when grown on BME + Col I, sh*B3galt6* cells failed to do so. These data suggest that B3GALT6 is required for both the viability of dormant tumor cells and their outgrowth.

We next performed analogous 3D assays using D2A1 cells. When grown on BME, a significant decrease in viable cell number was observed for sh*B3galt6* vs. sh*Scrambled* D2A1 cells at D12 (*p*=0.003), and sh*B3galt6* D2A1 cells failed to exhibit the characteristic spindle-like morphology adopted by sh*Scrambled* D2A1 cells that is associated with colony outgrowth (**Fig. S5C, D**). Notably, this same difference in phenotype was observed in sh*B3galt6* D2A1 cells (**Fig. S5E**) grown on BME + Col I. Moreover, a progressive decrease in viable cell numbers was observed for sh*B3galt6* D2A1 cells grown on BME + Col I at D4 (*p*=0.0014), D8 (*p*=0.0002), and D12 (*p*<0.0001) compared to control cells (**Fig. S5F**). These data indicate that, B3GALT6 is required for cell viability, as well as outgrowth, in proliferative D2A1 cells. Taken together, these findings suggest that B3GALT6 is a critical regulator of cell survival in dormant tumor cells, irrespective of whether dormancy is therapy-associated or induced by the microenvironment.

### Heparan sulfate synthesis is upregulated during dormancy and associated with poor outcomes in breast cancer patients

In light of our observations implicating B3GALT6-mediated proteoglycan biosynthesis as a key determinant of dormant tumor cell persistence in therapy-associated and microenvironment-induced models of dormancy, we next asked how proteoglycan synthesis is regulated during dormancy.

Because B3GALT6 catalyzes the synthesis of a linker that is common to, and required for, the production of both heparan sulfate and chondroitin sulfate (**Fig. 3A**), we assessed the biosynthesis of these GAGs under dormancy conditions. First, we applied heparan sulfate and chondroitin/dermatan sulfate KEGG mRNA biosynthesis signatures to IVD gene expression data derived from Her2-dependent cells. This revealed that enzymes involved in heparan sulfate synthesis and modification are upregulated (**Fig. S6A**) – whereas those involved in chondroitin/dermatan sulfate synthesis and modification are downregulated (**Fig. S6B**) – during dormancy in vitro.

Addition of the hexosamine residue *N*-Acetylglucosamine (GlcNAc) to the proteoglycan tetrasaccharide linker results in heparan sulfate synthesis, whereas addition of *N*-Acetylgalactosamine (GalNAc) to this linker results in chondroitin/dermatan sulfate synthesis^49^. Notably, the enzymes (*Extl2*, *Extl3*) that direct heparan sulfate synthesis by catalyzing the addition of GlcNAc to the tetrasaccharide linker are reversibly upregulated during dormancy in vitro (**Fig. S6C**); in contrast, the enzymes (*Csgalnact1*, *Csgalnact2*) that direct chondroitin sulfate synthesis by catalyzing the addition of GalNAc to this linker are reversibly downregulated during dormancy (**Fig. S6D**). Consistent with this, we identified the heparan sulfate polymerase *Ext2* as a hit in the CRISPR-Cas9 screen, whereby sgRNAs targeting *Ext2* were selected against during dormancy, as were sgRNAs targeting *B3galt6* (**Fig. 2D**). In contrast, negative selection for the chondroitin sulfate polymerase *Chpf* was not observed in our screen (**Fig. S2C**). These associations suggest the preferential synthesis of heparan sulfate rather than chondroitin/dermatan sulfate in dormant RTCs.

To quantify levels of heparan sulfate and chondroitin sulfate GAGs as a function of dormancy, we performed liquid chromatography-mass spectrometry (LC/MS) on cell lysates isolated at baseline (D0, proliferative) or post-doxycycline withdrawal (D7, dormant). Heparan sulfate was readily detectable in Her2-dependent tumor cells at D0 and its levels were significantly higher in dormant tumor cells at D7 (*p*<0.0001) (**Fig. 6A**). In contrast, chondroitin sulfate levels were >100-times lower than heparan sulfate at baseline (D0) and did not change significantly during dormancy (**Fig. 6A**).

**Fig. 6:**
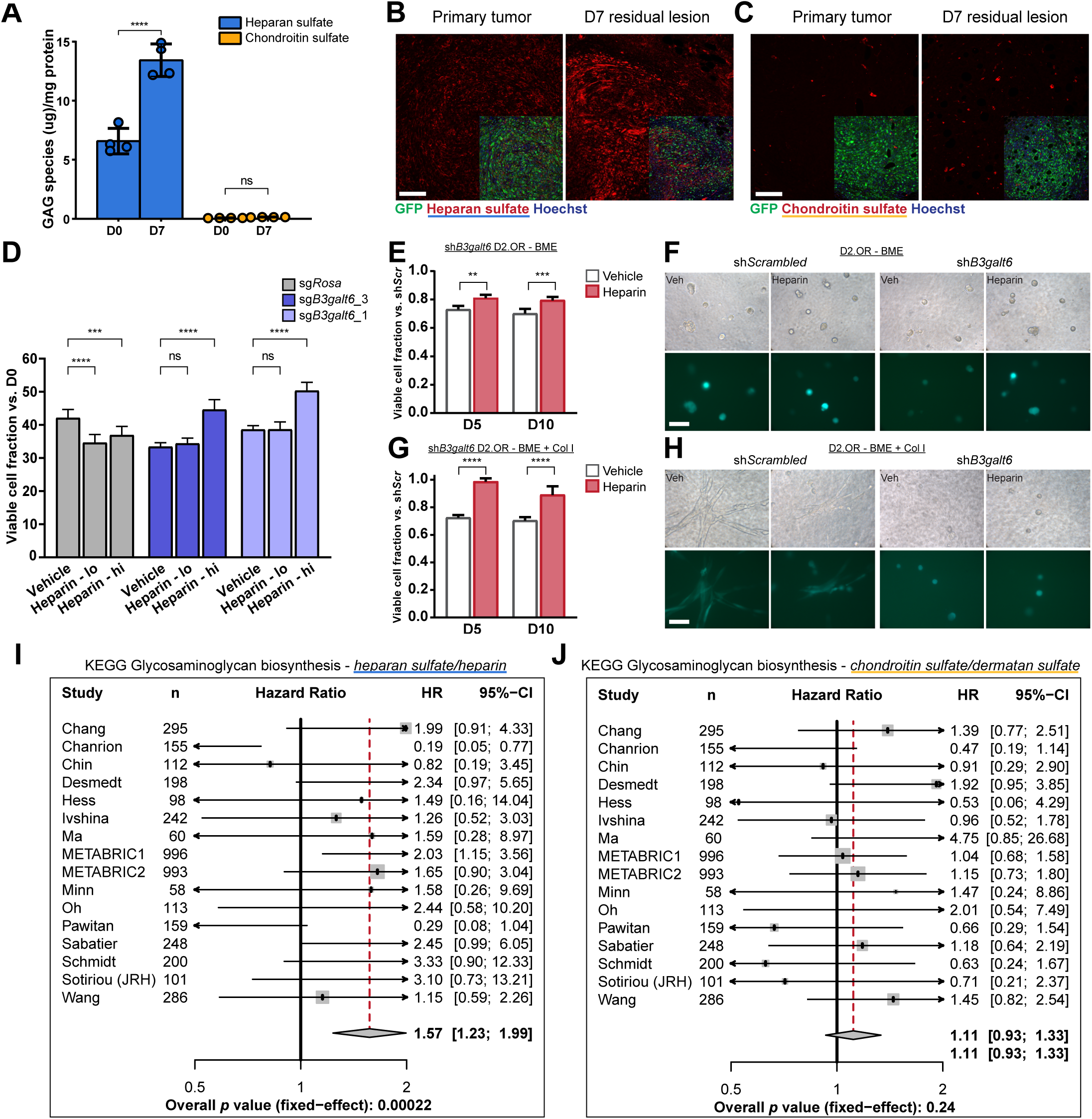
Heparan sulfate synthesis is upregulated during dormancy and associated with worse recurrence-free survival in breast cancer patients. **A.** Liquid chromatography/mass spectrometry (LC/MS) analysis of heparan sulfate (blue) and chondroitin sulfate (yellow) GAGs performed on D0 and D7 in vitro samples. Data are represented as mean ± SD. ns=non-significant, \*\*\*\**p*<0.0001. **B.** Immunofluorescence for heparan sulfate or **C.** chondroitin sulfate (right panels) (red) in PTs and D7 RLs derived from Her2-dependent-Cas9 cells with sg*Rosa* (green). Scale bar=100µm. **D.** Viable cell numbers measured at D7 following daily treatment of Her2-dependent cells with vehicle, heparin low dose (lo; 5µg/ml), or heparin high dose (hi; 25µg/ml). ns=non-significant, \*\*\**p*<0.001, \*\*\*\**p*<0.001. **E.** Viable shB3galt6 cell numbers normalized to shScrambled controls measured at D5 and D10 following treatment of D2.OR cells in BME with vehicle (grey) or heparin (25µg/ml, red) and **F.** fluorescence images. \*\**p*<0.01, \*\*\**p*<0.001. Scale bar=100µm. **G.** Viable shB3galt6 cell numbers normalized to shScrambled controls measured at D5 and D10 following treatment of D2.OR cells in BME + Col I with vehicle (grey) or heparin (25µg/ml, red) and **H.** fluorescence images. Scale bar=100µm. ****p<0.0001. **I.** Forest plots of hazard ratios (HR) and 95% confidence intervals (CI) as a function of heparan sulfate/heparin or **J.** chondroitin sulfate/dermatan sulfate KEGG biosynthesis signatures in breast cancer patients recurring within 10 years after initial treatment. Red dashed lines depict the shift in HR across 16 human datasets.

We next evaluated GAG levels in vivo as a function of dormancy using heparan or chondroitin sulfate-specific antibodies. We first confirmed the specificity of these antibodies by demonstrating that pre-treatment of tissue sections with heparin lyase or chondroitinase eliminated the signal detected for heparan sulfate or chondroitin sulfate, respectively (**Fig. S6E, F**). Immunofluorescence staining revealed that Her2-dependent PTCs (green) expressed heparan sulfate but not chondroitin sulfate (**Fig. 6B, C**; **S6E, F**). Moreover, heparan sulfate levels increased in dormant tumor cells at D7, whereas chondroitin sulfate remained undetectable (**Fig. 6B, C**). Thus, both LC/MS and immunofluorescence data demonstrate an increase in heparan sulfate abundance during dormancy, both in vitro and in vivo.

Defects in differentiation resulting from heparan sulfate impairment in myoblasts^50^ and mouse embryonic stem cells^51^ in vitro have been reported to be partially rescued by the addition of exogenous heparin, a highly sulfated heparan sulfate variant. Therefore, we performed an analogous experiment to determine whether heparin addition could rescue the impaired cell survival observed in B3GALT6-depleted cells, as would be predicted if heparan sulfate promotes the survival of dormant tumor cells.

We harvested Her2-dependent sg*Rosa* and sg*B3galt6* cells at baseline (D0, proliferative) and in cells treated with vehicle, low-dose heparin, or high-dose heparin for 7 days post-doxycycline withdrawal (D7, dormant). As above, sg*B3galt6* cells exhibited decreased survival at D7 compared to control cells **(Fig. 6D)**. Consistent with reports that heparin supplementation in heparan sulfate-replete cells may dampen signaling by competing with endogenous heparan sulfate^52^, we observed that both low- and high-dose heparin treatment in sg*Rosa* cells modestly decreased viable RTC number vs. vehicle-treated controls (**Fig. 6D**). In contrast, exogenous addition of high-dose heparin to sg*B3galt6* RTCs markedly increased the number of viable tumor cells at D7 vs. vehicle-treated controls (*p*<0.0001) (**Fig. 6D**). These data indicate that heparin supplementation is sufficient to rescue the survival defect induced by loss of B3GALT6 in dormant RTCs.

To extend these findings to a microenvironment-induced model of dormancy, we plated D2.OR cells transduced with sh*B3galt6* or control sh*Scrambled* hairpins in 3D in the presence of vehicle or heparin. As before, loss of B3GALT6 resulted in a pronounced survival defect in dormant D2.OR cells grown on BME (**Fig. 6D**). Notably, addition of exogenous heparin partially rescued the decrease in viable cell number observed in dormant D2.OR sh*B3galt6* cells vs. sh*Scrambled* at D5 (*p*=0.003) and D10 (*p*=0.0009) (**Fig. 6E, F**). Additionally, when D2.OR cells were plated under outgrowth conditions (BME + Col I), heparin potently rescued the decrease in cell numbers observed in dormant D2.OR sh*B3galt6* cells vs. sh*Scrambled* at D5 (*p*<0.0001) and D10 (*p*<0.0001) (**Fig. 6G**). These findings are concordant with those observed in Her2-dependent cells and consistent with the hypothesis that heparan sulfate GAGs are required for maintaining dormant RTC survival and subsequent outgrowth. However, heparin failed to rescue the rounded morphology associated with sh*B3galt6*, suggesting that D2.OR cells may transduce the Collagen I signal to stimulate the spindle-like morphology of outgrowths independently of heparin/heparan sulfate. (**Fig. 6H**).

To this point, our functional data in preclinical models suggested a role for heparan sulfate, but not chondroitin sulfate, in dormant tumor cell survival and we had observed the concordant dormancy-associated upregulation of a heparan sulfate KEGG mRNA biosynthesis signature, expression of rate-limiting enzymes for heparan sulfate synthesis, and levels of heparan sulfate, as well as a lack of dormancy-associated changes in these same features for chondroitin sulfate. Therefore, we applied the KEGG heparan sulfate and chondroitin/dermatan sulfate mRNA biosynthetic signatures to gene expression data from primary breast cancers in ∼4400 patients with known recurrence outcomes^38^. Strikingly, and consistent with a role for heparan sulfate in enhancing tumor cell fitness during dormancy and recurrence, the expression of heparan sulfate biosynthetic enzymes in early stage primary breast cancers was strongly associated with poorer recurrence-free survival (overall *p* value=2.2e-04) (**Fig. 6I**). In contrast, no association was observed between recurrence-free survival and expression chondroitin/dermatan sulfate biosynthetic enzymes (**Fig. 6J**). These data provide further evidence specifically implicating heparan sulfate, rather than chondroitin sulfate, biosynthesis in promoting breast cancer recurrence in patients.

Our findings demonstrate that enzymes involved in heparan sulfate synthesis, as well as total heparan sulfate levels, are upregulated in dormant tumor cells both in vivo and in vitro and that heparin supplementation partially rescues the survival defect induced by B3GALT6 loss in RTCs both in the context of therapy-associated dormancy and microenvironment-induced dormancy. In contrast, we found that chondroitin sulfate levels are orders of magnitude lower than heparan sulfate in tumor cells and are not upregulated during dormancy. Consistent with this, the enzymes involved in chondroitin sulfate synthesis are downregulated during dormancy. Taken together with our other findings, these data suggest a selective and essential role for heparan sulfate in mediating dormant RTC survival.

### Heparan sulfate 6-*O*-sulfation is selectively upregulated during dormancy and potentiates FGF1 signaling

Having identified a key role for B3GALT6-mediated heparan sulfate biosynthesis in promoting RTC survival, we asked if heparan sulfate undergoes dynamic modifications during dormancy, as was suggested by the reversible upregulation of gene expression for modifying enzymes, such as sulfotransferases (*e.g.*, *Ndst2*, *Hs2st1*, *Hs6st1*), within the heparan sulfate signature (**Fig. S6C**). Accordingly, we examined the extent and sites of heparan sulfation since this modification is a key determinant of the ligand binding properties of GAG side chains^23,53^.

To accomplish this, we isolated cell lysates from Her2-dependent cells grown in vitro at D0 (*baseline*), D7 and D28 (*deinduction*), and D28+ (*reinduction*) and performed glycan reductive isotope labeling followed by LC/MS (GRIL-LC/MS)^54^ (**Fig. S7A**). This revealed a significant and progressive increase in the average sulfation level per heparan sulfate-derived disaccharide at D7 and D28 of dormancy vs. D0 baseline (*p*<0.0001), which remained elevated after 72 hr of Her2 reinduction (**Fig. S7B**). This dormancy-associated increase in sulfation indicates that, in addition to increased overall levels of heparan sulfate GAGs during dormancy, the extent to which these GAGs are sulfated is also dynamically upregulated in RTCs.

Next, we wished to determine whether dormancy-selective patterns of heparan sulfation were present. Of the 3 (out of 4 total) sulfation sites on heparan sulfate-derived disaccharides that we interrogated (**Fig. 7A-C; S7C**), 6-*O*-sulfation exhibited the most dramatic fold-increase in abundance during dormancy **(Fig. 7A-D)**. We also examined the relative abundance of specific disaccharide motifs^55^ as a function of dormancy (**Fig. S7D, E**). Only D2S6 and D0A6 disaccharides showed upregulation during dormancy in a manner that was reversible following exit of cells from the dormant state (**Fig. S7F, G**). This indicates that 6-*O*-sulfation on glucosamine tends to co-occur in non-sulfated uronic acid-*N*-acetylated glucosamine (D0A6) or 2-*O*-sulfated uronic acid-*N*-sulfated glucosamine (D2S6) disaccharides.

**Fig. 7:**
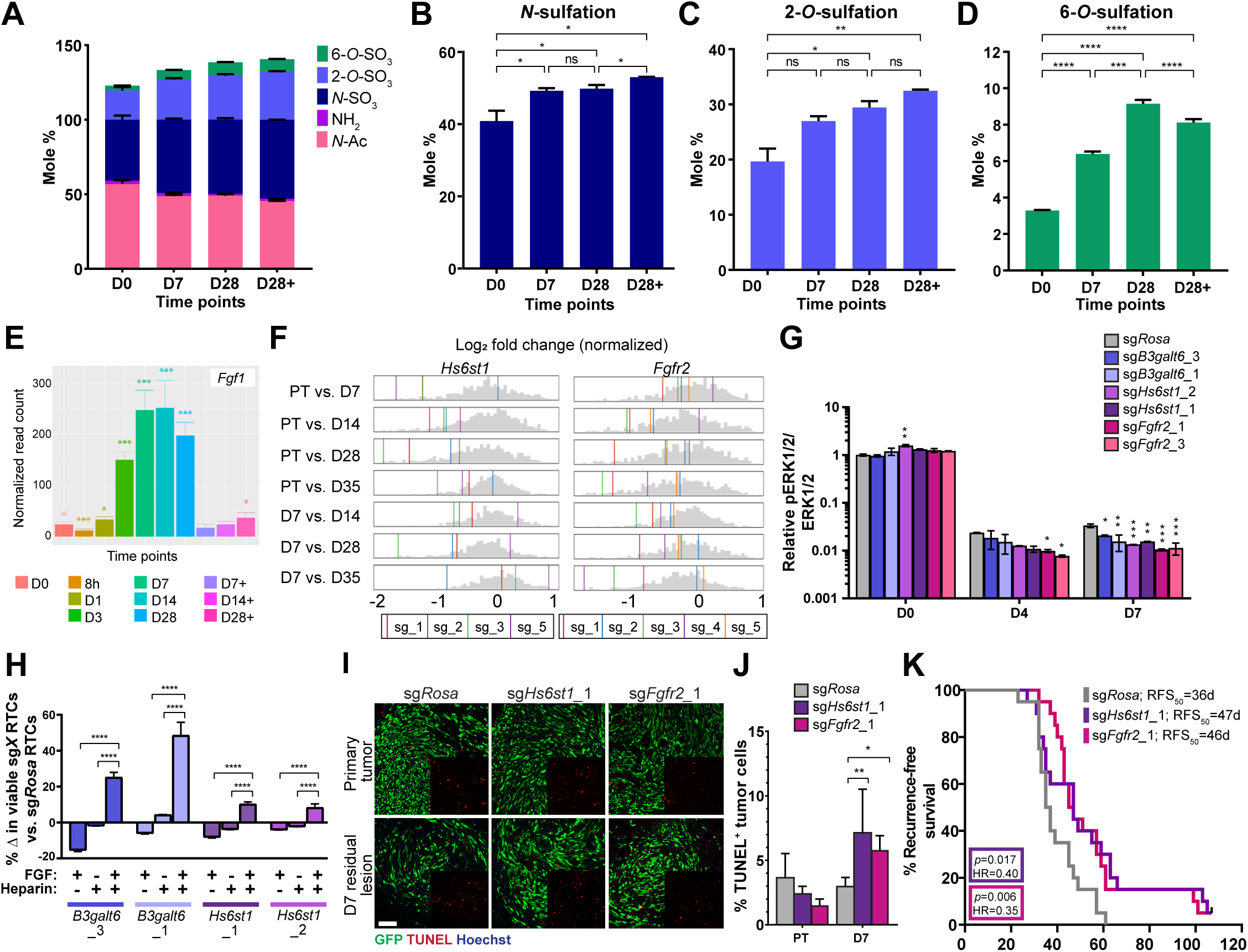
Heparan sulfate 6-*O*-sulfation is selectively upregulated during dormancy and promotes RTC survival by potentiating FGF1 signaling. **A.** LC/MS on D0 (*baseline*), D7, D28 (*deinduction*), and D28+ (*reinduction*) IVD samples identifying the molar percentages of modifications on disaccharide units comprising iduronic/glucuronic acid and glucosamine. Relative proportions of N-acetylglucos-amine (N-Ac; light pink), unsubstituted glucosamine (NH_2_; dark pink), **B.** N-sulfoglucosamine (N-S; dark blue), **C.** Uronyl-2-*O*-sulfates (2-*O*-SO_3_; light blue), and **D.** Glucosaminyl-6-*O*-sulfates (6-*O*-SO_3_; green) are depicted as mean ± SD. ns=non-significant, \**p*<0.05, \*\**p*<0.01, \*\*\**p*<0.001, \*\*\*\**p*<0.001. **E.** Normalized read counts indicating reversible upregulation of *Fgf1* at 8h, D1, D3, D7, D14, D28 (*deinduction*) time points. Asterisks indicate significant changes in normalized read counts vs. D0 (*baseline*) \**p*<0.05, \*\*\**p*<0.001. **F.** Log_2_ fold-change (normalized) gene-level effect size plots for *B3galt6* in pairwise comparisons using PT or D7 as baselines. Grey histogram depicts the background distribution of sgRNAs in the CRISPR-Cas9 screen, colored vertical lines depict each sgRNA targeting *Hs6st1* or *Fgfr2.* **G.** Western blot analysis of phospho-ERK1/2 (pERK1/2) and total ERK1/2 (tERK1/2) levels in sg*Rosa*, sg*B3galt6*_3, sg*B3galt6*_1, sg*Hs6st1*_2, sg*Hs6st1*_1, sg*Fgfr2*_2, and sg*Fgfr2*_3 Her2-dependent-Cas9 cells at D0 (*baseline*), D4, and D7 (*deinduction*) time points. Numbers indicate relative quantification of pERK1/2/tER- K1/2 signal normalized to sgRosa levels within each time point. **H.** Viable RTC counts at D7 quantifying the percentage change relative to sgRosa RTCs in sg*B3galt6*_3, sg*B3galt6*_1, sg*Hs6st1*_2, and sg*Hs6st1*_1 Her2-dependent-Cas9 cells treated with FGF1 (25ng/ml), heparin (25µg/ml), or FGF1 (25ng/ml) + heparin (25µg/ml). \*\*\*\**p*<0.0001. **I.** Immunofluorescence and **J.** quantification for TUNEL (red) in Her2-dependent-Cas9 sg*Rosa*, sg*Hs6st1*_1, and sg*Fgfr2*_1 (green) PTs and D7 RLs. Scale bar=100µm. Quantification in the sg*Rosa* (grey), sg*Hs6st1*_1 (purple), and sg*Fgfr2*_1 (pink) groups is represented as mean ± SD. \**p*<0.05, \*\**p*<0.01. **K.** Kaplan-Meier analysis of recurrence-free survival for the sg*Rosa* (grey), sg*Hs6st1*_1 (purple), and sgF*g- fr2*_1 (pink) groups. RFS_50_=median time-to-recurrence.

Heparan 6-*O*-sulfation has been reported to promote FGF signaling by increasing the binding affinity of some FGF ligands for heparan sulfate in the ternary heparan sulfate-FGF-FGFR complex that is required for FGF signaling^56,57^ (**Fig. S7H**). To identify which FGF ligands are expressed in dormant tumor cells in an abundant, but reversible, manner we interrogated the expression of the 15 paracrine FGFs that require heparin/heparan sulfate as a co-factor for signaling^58^. This revealed that FGF1 is dramatically and persistently upregulated (>10-fold) during dormancy as early as 3 days following doxycycline withdrawal, and that its expression is potently suppressed within 48 hr of doxycycline re-addition (D7+, D14+, D28+) (**Fig. 7E**). Other notable FGF ligands included FGF2, which displayed only minor upregulation during dormancy that was not reversible, and FGF7, which displayed reversibility but was expressed at low levels with only modest dormancy-associated upregulation (**Fig. S7I**).

Our finding that FGF1 expression is markedly upregulated in dormant tumor cells was particularly striking in light of recent data identifying heparan 6-*O*-sulfation as a strong binding site for FGF1, but not FGF2^59^. This suggested the possibility that dormant tumor cells might be specifically dependent on heparan sulfate 6-*O*-sulfation as a consequence of its interaction with FGF1.

To test whether FGF1 regulates RTC survival, we determined the number of dormant RTCs that survived Her2 downregulation in vitro following the depletion of endogenous FGF1 using CRISPR-Cas9-based deletion. After confirming successful editing by *Fgf1* sgRNAs by ICE analysis (**Fig. S7J**), we performed an IVD assay using Her2-dependent*-*Cas9 cells transduced with sg*Fgf1* or sg*Rosa*. Deletion of FGF1 further accentuated the decrease in viable RTC numbers that occurs at D7 (*deinduction*) vs. D0 (*baseline*) when compared to the sg*Rosa* controls (**Fig. S7K**). This suggests that tumor cell-autonomous FGF1 signaling promotes dormant RTC survival.

To determine whether FGF acts via heparan 6-*O*-sulfation to mediate B3GALT6-associated RTC survival, we examined the expression of the three sulfotransferases, *Hs6st1-3,* that catalyze 6-*O*-sulfation (**Fig. S7H**), particularly those on the D2S6 and D0A6 6-*O*-sulfated disaccharide motifs^53,56^ identified above. Of these, only *Hs6st1* was found to be abundantly expressed in Her2-dependent cells and only *Hs6st1* was reversibly upregulated during dormancy (**Fig. S7L**). Notably, *Hs6st1* was identified as a putative pro-survival hit in our CRISPR-Cas9 screen, which exhibited increasingly apparent effects at later dormancy time points (**Fig. 2D, 7F**).

We also evaluated the expression of the four FGFRs, *Fgfr1-4*, that participate in FGF signaling. Of these, *Fgfr1* and *Fgfr2* are abundantly expressed in Her2-dependent cells and are each reversibly upregulated during dormancy (**Fig. S7L**); *Fgfr2* was upregulated ∼4-fold during dormancy, whereas *Fgfr1* was upregulated ∼1.5-fold. Although FGF1 can signal through both FGFR1 and FGFR2, *Fgfr2* was identified as a putative pro-survival hit in our CRISPR-Cas9 screen, whereas sgRNAs targeting *Fgfr1* displayed no selection (**Fig. 2D, S2C, 7F**). Accordingly, we hypothesized that 6-*O*-sulfated heparan sulfate and FGFR2 may act together to promote FGF1 signaling and RTC survival, and prioritized studies of FGFR2 as a candidate for maintaining dormant RTC survival.

To determine whether heparan 6-*O*-sulfation and FGFR2 act in concert to promote FGF1 signaling and dormant RTC survival, we first sought to evaluate FGF activity in dormant tumor cells in vitro, as well as the effect of inhibiting heparan sulfate biosynthesis or 6-*O*-sulfation on FGF signaling.

After confirming successful editing by *Hs6st1* and *Fgfr2* sgRNAs by ICE analysis (**Fig. S7M**), we performed an IVD assay using Her2-dependent-Cas9 cells transduced with sg*B3galt6*, sg*Hs6st1*, sg*Fgfr2*, or sg*Rosa*. We reasoned that if increased heparan sulfate synthesis and 6-*O*-sulfation by dormant RTCs enhances endogenous FGF signaling, then impairing heparan sulfate synthesis (sg*B3galt6*), or 6-*O*-sulfation (sg*Hs6st1*), should attenuate FGF signaling (**Fig. S7N**). Furthermore, we predicted that if enhanced FGF signaling in dormant cells is mediated by FGFR2, depleting *Fgfr2* should also attenuate FGF signaling during dormancy.

To test this hypothesis, we assessed levels of activated ERK1/2^60^ in each of the above genetic contexts, using two sgRNAs each for sg*B3galt6*, sg*Hs6st1*, and sg*Fgfr2*. As anticipated, pERK1/2 levels normalized to total ERK1/2 were >10-fold higher in Her2-dependent proliferating cells at D0 (*baseline*) compared to D4 or D7 dormancy time points (**Fig. 7G, S7O**). Moreover, pERK1/2:ERK1/2 levels in the presence of Her2 expression (D0) were largely unaffected by deletion of *B3galt6*, *Hs6st1*, or *Fgfr2*. In contrast, pERK1/2:ERK1/2 levels were significantly and progressively diminished in sg*B3galt6*, sg*Hs6st1*, and sg*Fgfr2* cells compared to *sgRosa* controls following Her2 downregulation (*deinduction*) induced by doxycycline withdrawal (**Fig. 7G, S7O**). This additional impairment of pERK1/2:ERK1/2 levels caused by deletion of components of the FGF signaling pathway that are expressed in RTCs in a dormancy-specific manner is consistent with a model in which endogenous FGF signaling is active during dormancy, and requires B3GALT6, HS6ST1, and FGFR2 for its maintenance.

### Upregulation of heparan sulfate 6-*O*-sulfation during dormancy promotes RTC survival and recurrence by potentiating FGF signaling

Given the above findings, we assessed whether the addition of exogenous FGF ligands can increase the number of viable RTCs in vitro. Indeed, we found that addition of FGF1 to sg*Rosa* cells for 4 days initiated concurrently with doxycycline withdrawal resulted in a near doubling of the number of viable RTCs compared to vehicle controls (dotted line) (**Fig. S7P**). This FGF1-induced increase in viable dormant RTCs was impaired in sg*B3galt6*, sg*Hs6st1*, and sg*Fgfr2* cells vs. sg*Rosa* controls (**Fig. S7P**). These findings suggest that heparan sulfate (synthesized by B3GALT6), particularly 6-*O*-sulfation (catalyzed by HS6ST1), as well as FGFR2 are key components of FGF signal transduction in dormant RTCs.

Next, we asked if the impaired response to FGF1 in sg*B3galt6* and sg*Hs6st1* RTCs compared to sg*Rosa* RTCs could be rescued by the addition of heparin, which is natively sulfated, including at the 6-*O* site. We treated Her2-dependent tumor cells with FGF1, heparin, or a combination of FGF1 + heparin for 7 days post-doxycycline withdrawal (D7, dormant). We found that heparin could rescue the blunted response to exogenous FGF1 stimulation observed in sg*B3galt6* and sg*Hs6st1* cells (**Fig. 7H**). These data are consistent with a model wherein heparan 6-*O-*sulfation potentiates FGF signaling to promote the survival of dormant RTCs in vitro.

We wished to determine whether heparan 6-*O-*sulfation and FGF signaling promote dormant RTC viability and tumor recurrence as suggested by the CRISPR screen results (**Fig. 2D, 7F**). We assessed viable RTC number for sg*Rosa*, sg*B3galt6*, sg*Hs6st1*, and sg*Fgfr2*-transduced Her2-dependent-Cas9 cells at D7 (*deinduction*). Consistent with our hypothesis, dormant tumor cells deleted for each of these genes exhibited significantly impaired survival during dormancy vs. sg*Rosa* control cells (**Fig. S7Q**). This suggests that heparan 6-*O-*sulfation potentiates endogenous FGF signaling mediated by FGFR2 to promote the survival of dormant RTCs in vitro.

To extend these findings in vivo, we injected GFP-labeled sg*Rosa*, sg*Hs6st1*, and sg*Fgfr2* Her2-dependent-Cas9 cells into mice maintained on doxycycline. Her2-driven PT samples (*baseline*) as well as D7 RL samples following doxycycline withdrawal were harvested for TUNEL staining. This revealed a significant increase in the percentage of TUNEL+ sg*Hs6st1* (*p*=0.003) and sg*Fgfr2* (*p*=0.04) tumor cells in D7 RL vs. sg*Rosa* controls (**Fig. 7I, J**). In contrast, we found no significant differences in the percentage of TUNEL+ tumor cells in sg*Hs6st1,* sg*Fgfr2,* and *sgRosa* PT samples. These data indicate that sg*Hs6st1* and sg*Fgfr2* dormant RTCs display a selective impairment in RTC, but not PTC, survival compared to the controls, in a manner analogous to our findings for sg*B3galt6*.

Finally, to determine the impact of HS6ST1 and FGFR2-mediated pro-survival effects on tumor recurrence, we performed recurrence-free survival assays in mice bearing dormant RLs derived from sg*Hs6st1* or sg*Fgfr2* Her2-dependent tumor cells. Analogous to our findings for sg*B3galt6*, mice bearing dormant RLs containing sg*Hs6st1* or *Fgfr2* tumor cells exhibited delayed recurrence-free survival compared to sg*Rosa* controls (*p*=0.02, HR=0.42; and *p*=0.006, HR=0.35, respectively) (**Fig. 7K**). This demonstrates that HS6ST1 and FGFR2 are each required for efficient tumor recurrence.

In aggregate, the above studies are consistent with a model in which HS6ST1 is reversibly upregulated during dormancy, which results in the selective upregulation of heparan sulfate 6-*O*-sulfation in dormant RTCs and potentiation of FGF1 signaling via FGFR2, thereby enhancing dormant RTC survival. This, in turn, promotes tumor recurrence, as reflected by the delay in the median time-to-recurrence observed for tumor cells deleted for either *Hs6st1* or *Fgfr2*.

## Discussion

Residual tumor cells (RTCs) that persist following primary tumor therapy have long been recognized as the precursors of treatment-refractory recurrent disease that determines patient mortality. While a small number of post-adjuvant clinical trials to identify and target dormant RTCs in early stage breast cancer patients are currently underway^61,62^, there is a pressing need to identify unique biological vulnerabilities of dormant RTCs.

In this study, we address this unmet clinical need by identifying novel, actionable dependencies of RTC fitness that could be leveraged to prevent tumor recurrence. Utilizing a combination of in vivo and in vitro derived gene expression data from mouse models and breast cancer patients, a CRISPR-Cas9-based screen, and functional validation studies in mouse models for tumor dormancy and recurrence, we found that dormant RTCs create – in a cell autonomous manner – an extracellular environment that is conducive to their own survival. In particular, we determined that dormant RTCs selectively upregulate the B3GALT6-mediated synthesis of heparan sulfate proteoglycans in both therapy-associated and microenvironment-induced models of dormancy. Underscoring the clinical relevance of these data, we observed that increased heparan sulfate synthesis in primary tumors in patients is associated with poor recurrence-free survival. Mechanistically, we found that dormant RTCs reversibly upregulate not only B3GALT6-mediated heparan sulfate synthesis, but also HS6ST1-mediated 6-*O*-sulfation of heparan sulfate proteoglycans, as well as *Fgf1* and *Fgfr2* expression, in a dormancy-specific manner. These orchestrated effects result in enhanced FGF1 signaling via FGFR2, enhanced dormant RTC survival, and accelerated tumor recurrence. In aggregate, our findings identify the B3GALT6-heparan sulfate/HS6ST1-6-*O*-sulfation/FGF1-FGFR2 signaling axis (**Fig. S7N**) as a potential therapeutic target for preventing tumor recurrence from dormant RTCs.

Proteoglycans have pleiotropic functions in both normal physiology and during cancer initiation and progression^23^. Individual proteoglycans have been implicated as regulators of tumor cell growth, angiogenesis, epithelial-to-mesenchymal transition, anoikis, extravasation, and colonization of metastatic locations^63^. In contrast, little is known about potential roles for proteoglycans during the dormant phase of cancer progression^64^, and fewer studies still have dissected the function of the protein core of proteoglycans from that of their GAG chains.

Our findings identify B3GALT6 as a potent regulator of dormant RTC survival and recurrence. To date, a role for B3GALT6 in cancer has not been reported. Because B3GALT6 is essential for synthesis of the tetrasaccharide required for covalent linkage of GAG chains to their protein core^28^, genetic deletion of *B3galt6* permits the role of GAGs in tumor progression to be specifically investigated, as opposed to their protein cores. Prior studies of B3GALT6 in vertebrate models have been limited to genetic loss in a zebrafish model^46^ generated for study of two pathogenic conditions observed in patients with biallelic loss of *B3GALT6*: spEDS and SEMD-JL1^29,31^. Accordingly, while zebrafish with *b3galt6* loss-of-function recapitulate the connective tissue defects observed in spEDS and SEMD-JL1 patients, the extent to which the pathways underlying B3GALT6-mediated functions in connective tissues, and those explaining the dependence of dormant epithelial tumor cells on this enzyme, are shared – if at all – remains to be determined.

The levels and spatial patterns of heparan sulfation strongly influence the extracellular ligand milieu and the extent to which proteoglycans elicit signaling in cells^26^. In addition to the dormancy-associated upregulation of heparan sulfate proteoglycans in our model, we discovered an upregulation of the extent of heparan sulfation in dormant residual tumor cells. A recent study in melanoma cells identified increased levels of a transporter of the sulfate donor PAPS (*i.e.*, SLC35B2) and downstream heparan sulfation as a dependency of melanoma cells that are refractory to MAPK pathway inhibition^65^. In light of our study, these data may suggest the possibility that increased heparan sulfation may constitute an adaptive mechanism of drug resistance in multiple cancer types, although the site of heparan sulfation and dependence of this phenotype on FGF signaling were not addressed in this study.

Heparan 6-*O* sulfation is required for formation of the ternary heparan sulfate-FGF-FGFR complex and, therefore, FGF pathway activity^56,57^. In particular, 6-*O*-sulfation is a strong determinant of FGF1 binding^59^. FGF signaling is classically thought to play a mitogenic function in tumor cells^8,66^. However, recent data suggest a context-dependent role for FGF signaling in promoting breast cancer therapy resistance and growth arrest. FGF2 derived from osteogenic cells in the bone marrow can suppress the expression of estrogen receptor (ER) in disseminated breast cancer cells via FGFR1, rendering them resistant to endocrine therapy^67^. Furthermore, exogenously-derived FGF2 induces cell cycle arrest in ER+ breast cancer cell lines, such as MCF7 and T47D^68^, and can induce the expression of the pro-dormancy transcription factor ZFP281 in early disseminated cancer cells^69^. In contrast to these examples in which FGF2 can promote dormancy predominantly via FGFR1, our data suggest dormant RTC survival is mediated by tumor cell-autonomous FGF1 in a heparan sulfate-FGFR2 dependent manner. In this regard, we found that genetic depletion of FGFR2 induced RTC apoptosis and significantly delayed recurrence. Consistent with this, our CRISPR screen identified *Fgfr2*, but not *Fgfr1*, as a vulnerability in dormant RTCs. These data suggest the intriguing possibility that highly selective FGFR2 pharmacological inhibitors currently in clinical trials in other solid tumors (NCT04526106^70^, NCT04071184^71^) could be leveraged in the adjuvant or post-adjuvant setting, to exploit the dependency of dormant RTCs on FGF signaling for their survival and recurrence.

Although heparan 6-*O*-sulfation has been reported to increase the affinity of VEGFA binding to VEGFR2^72^, both VEGFA and VEGFR2 gene expression levels are downregulated during dormancy in our model, thereby providing further evidence for the selective effects of FGF signaling in the context of dormancy and heparan 6-*O*-sulfation. Consistent with this, we have not observed evidence supporting angiogenic impairment as a mechanism of dormancy in our model^6^. Notably, desulfation of heparan sulfate at the 6-*O* position is catalyzed by the sulfatases, SULF1 and SULF2^53^. As such, our data demonstrating that heparan 6-*O*-sulfation is preferentially increased during dormancy is compatible not only with models in which sulfotransferase activity is upregulated, as we have shown occurs during dormancy, but also with models in which sulfatase activity is downregulated. Although we observe no downregulation of *Sulf1* and *Sulf2* levels in dormancy, further investigation into the regulation of 6-*O*-sulfation during breast cancer progression is a promising avenue of future research.

Intriguingly, our studies implicate heparan sulfate, rather than chondroitin sulfate, as the key GAG responsible for promoting RTC fitness. Although our study is focused on the glycan component of proteoglycans, the core protein may also play an important role beyond serving as a scaffold for the attachment of GAGs. Indeed, sequences within the core protein can influence the stoichiometry^73^ and sulfation pattern of GAG chains^72^, with important biological consequences^74^. The heparan sulfate proteoglycan family orchestrates diverse physiological functions^26^ and includes transmembrane syndecans (SDC1-4), 6 glycophosphatidylinositol (GPI)-anchored glypicans (GPC1-6), as well as secreted perlecan, agrin, and collagen XVIII that are deposited in the extracellular matrix^25^. Additionally, betaglycan (also known as TGFBR3) and a splice variant of CD44 (CD44v3) can be modified by GAGs, including heparan sulfate, and act as ‘part-time’ proteoglycans^25^. Determining which heparan sulfate proteoglycans contribute to RTC survival in breast cancer models, and whether their functions are dependent on the heparan sulfate GAG are intriguing directions to pursue.

In conclusion, our study identifies a dormancy-specific function for the B3GALT6-heparan sulfate/HS6ST1-6-*O*-sulfation/FGF1-FGFR2 signaling axis in maintaining RTC survival. In addition to ascribing a critical role to an understudied class of compounds, namely glycosaminoglycans, in breast cancer progression, our study leverages this information to identify novel therapeutic vulnerabilities for depleting RTCs in an effort to mitigate the development of incurable recurrences.

## Acknowledgments

We thank Dr. Junwei Shi and his laboratory for their advice on developing and performing the CRISPR-Cas9 screen, Dr. Mikala Egeblad for the gift of D2.OR and D2A1 cells, Jianping Wang for histology assistance, Mousumi Paulchakrabarti for GRIL-LC/MS assistance, and Cuyler Luck as well as members of the Chodosh laboratory for their invaluable input that led to the development of techniques used in this study. This project was supported in part by grants from the NIH and the Breast Cancer Research Foundation to L.A.C. and by a grant from DOD to A.S.

## Author contributions

Conceptualization, A.S., J.D.E., and L.A.C.; Methodology, A.S., M.L., B.C., T.P., D.K.P., C.J.S., G.K.B., T.T., B.B., M.E., and F.E.M.; Investigation, A.S., M.L., B.C., T.P., and D.K.P.; Writing, A.S. and L.A.C.; Funding Acquisition, A.S. and L.A.C.

## STAR Methods

### In vitro assays

*MTB/TAN*-derived primary tumor cells were cultured in DMEM (Corning, Cat. # 10-017-CV) containing 10% super calf serum (GeminiBio, Cat. # 100-510), 1% Penicillin/Streptomycin (Thermo Fisher Scientific, Cat. # 15-140-122), 1% Glutamine (Thermo Fisher Scientific, Cat. # 25030081), 2mg/ml Doxycycline (RPI, Cat. # D43020-250.0), 5mg/ml Prolactin (NHPP, Cat. # NIDDK-oPRL-21), 5mg/ml Insulin (GeminiBio, Cat. # 700-112P), 10ug/ml EGF (Millipore, Cat. # E4127), 1mg/ml Hydrocortisone (Sigma, Cat. # H0396), 1mM Progesterone (Sigma, Cat. # P7556).

For in vitro dormancy experiments, cells were plated on D-3 in medium as above, transitioned to medium as above containing 1% super calf serum on D-2, and transitioned to medium containing 1% super calf serum and no Doxycycline on D0. Plates were harvested at dormancy deinduction time points as desired, while medium containing 1% super calf serum and no Doxycycline was replaced on remaining plates weekly. Reinduction plates were harvested at time points as desired following 48h or 72h treatment with medium containing 1% super calf serum and doxycycline as indicated in the figures.

D2.OR and D2A1 cells were a gift from Dr. Mikala Egeblad at Cold Spring Harbor Laboratories. These cells were maintained in in DMEM (Corning, Cat. # 10-017-CV) containing 10% fetal bovine serum (GeminiBio, Cat. # 100-106), 1% Penicillin/Streptomycin (Thermo Fisher Scientific, Cat. # 15-140-122).

Cell viability assays (2D and 3D) were performed as per manufacturer’s instructions using the Cell Titer 96 Non-Radioactive Cell Proliferation Assay (Promega, Cat. # G4000) in 96-well plates.

### 3D assays

D2.OR and D2A1 cells were plated on top of a matrix of basement membrane extract (Cultrex 3D Basement Membrane Extract, Reduced Growth Factor, Trevigen Cat. # 3445-005-01) or a 1:1 ratio of basement membrane extract and neutralized type I collagen (Cultrex 3D Culture Matrix Rat Collagen I, Trevigen Cat. # 3447-020-01). For experiments in 96-well plates (Corning, Cat. # 353219), 50µl of the matrix was added per well and were solidified for 1h at 37°C. D2.OR and D2A1 cells were resuspended at 20,000 cells/ml in DMEM low glucose, low pyruvate medium (Thermo Fisher Scientific, Cat. # 11885092) containing 2% fetal bovine serum (GeminiBio, Cat. # 100-106), 2% basement membrane extract, 1% Penicillin/Streptomycin (Thermo Fisher Scientific, Cat. # 15-140-122), and 100µl per well was plated on top of the solidified matrices. Plates were harvested at desired time points and medium was replaced on remaining plates every 4 days. Cell viability in the harvested plates was assayed using the CellTiter Non-Radioactive Cell Proliferation Assay (Promega, Cat. # G4000) as per manufacturer’s instructions.

D2.OR and D2A1 cells were similarly cultured in 18 well Glass Bottom chamber slides (ibidi, Cat. # 81817) for immunofluorescence with the following modifications - 20µl of the matrix was added per well and wells were solidified for 1h at 37°C following which 200 cells were added to each well.

### RNA sequencing

*MTB/TAN* cells were cultured in vitro under dormancy conditions as described. Samples were harvested at baseline (D0), deinduction (8 h, D1, D3, D7, D14, D28), and reinduction (D7+, D14+, and D28+) timepoints. RNA isolation was performed using the RNeasy Mini kit (Qiagen, Cat. # 74106) and sequencing libraries were prepared using the TruSeq Stranded mRNA for NeoPrep kit (Illumina, Cat. # NP-202-1001). Sequencing was performed using 75-bp paired-end NextSeq (Illumina, Cat. # 20024907).

The quality of raw reads was assessed using FASTQC. Sequenced reads were mapped to the mm10 *Mus musculus* reference genome using Spliced Transcripts Alignment to a Reference (STAR). Gene-level read counts were determined using featureCounts. Principal component analysis (PCA) was performed using the top 1000 most variable genes assessed by per-gene standard deviations and was used to exclude distinct outliers from the downstream analysis. Read counts across samples were normalized and differentially expressed genes were identified using DESeq2.

### Animal experiments

All animal studies were approved by the University of Pennsylvania Institutional Animal Care and Use Committee (IACUC). In vivo competition assays and recurrence-free survival assays were performed as previously described. Briefly, 1e06 cells (1:1 mixture of GFP^+^ and mCherry^+^ cells for competition assays or unmixed cells for recurrence-free survival assays and the CRISPR-Cas9 screen) were transplanted into the inguinal #4 mammary fat pads of *nu/nu* female mice (NCRNU-F, Taconic) and administered 2mg/ml doxycycline/5% sucrose via drinking water. Mice were de-induced by switching to regular drinking water once mammary tumors reached target size (5×5mm for competition assays; 3×3mm for recurrence-free survival assays and the CRISPR-Cas9 screen). Mice were palpated thrice a week and time-to-recurrence was assessed by Kaplan-Meier analysis. At the time of harvest, mice were administered 50mg/kg EdU (i.p.) 2h prior to euthanasia.

### Plasmids and lentiviral production

LentiV_Cas9_puro (Addgene, Cat. # 108100) was used to generate Cas9 expressing *MTB/TAN*-derived primary tumor cells. LRG2.1 (Addgene, Cat. # 108098), LRG (Addgene, Cat. # 65656), or LRmCherry2.1 (Addgene, Cat. # 108099) vector backbones were utilized for cloning sgRNAs for CRISPR-Cas9 studies. For each sgRNA, sense and antisense oligos were phosphorylated and annealed and then ligated into BsmB1-digested vector. Ligated vectors were transformed into the chemically competent Stbl3 bacteria (Thermo Fisher Scientific, Cat. # C737303). Successfully transformed bacterial clones were picked from Ampicillin selective plates and isolated DNA was sequenced using a U6 primer to confirm sgRNA incorporation.

Lentiviruses were generated in HEK293T cells using the TransIT-293 transfection reagent (Mirus, Cat. # MIR2700) to introduce packaging plasmids pMD2.G (Addgene, Cat. # 12259; 3µg) and psPAX2 (Addgene, Cat. # 12260; 6µg), and 9µg of the desired backbone containing the sgRNA of interest. sgRNA lentiviruses were titered by serial dilution in *MTB/TAN*-derived primary tumor cells using the fluorophore associated with the vector backbones as a readout by on the Attune NxT flow cytometry (Thermo Fisher).

### CRISPR-Cas9 screen

Of the genes that were selectively enriched during dormancy in vitro that were encompassed by extracellular matrix-associated gene ontology terms, 95 genes were selected for CRISPR-Cas9 screening in vivo. In addition, positive control sgRNAs targeting genes that are known to be functional during disease progression (pro-proliferative and pro-survival genes) and negative control sgRNAs that are non-targeting were also included in this library. For the 95 genes, 4-5 sgRNAs were designed for each gene resulting in a total of 509 sgRNAs. Additionally, a size-matched library of non-targeting sgRNAs was prepared to aid data analysis.

sgRNAs were designed to target conserved functional domains that display low computationally predicted off-target scores using the GUIDES tool (http://guides.sanjanalab.org/), preferentially picking sgRNAs that have an A/T nucleotide at the 17^th^ position of the sgRNA sequence. This approach maximizes functionally ‘null’ mutations and circumvents the need for subcloning the cells, thus maintaining their heterogeneity. These sgRNA oligo pools were cloned into the LRG expression vector following which the plasmid pool was amplified, purified, and packaged into lentiviruses. To ensure that *MTB/TAN*-Cas9 cells receive only one sgRNA/cell, the lentiviral library was titered using the GFP selectable marker by flow cytometry (Attune NxT Thermo Fisher) and transduced at an MOI=0.3. Finally, the transduced cells were sorted (MoFlo Astrios, Beckman Coulter Life Sciences) and expanded prior to transplantation in vivo. All steps were performed such that each sgRNA was represented in >1500 cells/sgRNA to maintain coverage and improve the robustness of downstream analyses.

All samples harvested from mouse primary tumors and residual lesions were microdissected under a stereoscope and homogenized for genomic DNA extraction using the Quick-DNA Midiprep Plus kit (Zymo Research, Cat. # D4075). sgRNA inserts were amplified by PCR using the Phusion Flash High Fidelity PCR Master Mix (Thermo Fisher, Cat. # F548), followed by barcode and adapter addition. The libraries were then pooled, mixed with 5% PhiX (Thermo Fisher, Cat. # FC-110-3001) and massively parallel sequenced using the MiSeq reagent kit v3 150 cycle (Illumina, Cat. # MS-102-3001) on the MiSeq Instrument (Illumina).

### Histology

When harvesting 3D assays for immunofluorescence, medium was aspirated and immediately fixed in 2% paraformaldehyde (Santa Cruz Biotechnology, Cat. # sc-281692) in PBS for 20 min at room temperature. Cells were then permeabilized in PBS containing 0.5% Triton X-100 for 10 min at room temperature. Wells were then rinsed three times in 1x PBS containing 100mM glycine before proceeding with the immunofluorescence protocol.

For in vitro immunofluorescence/labeling studies, *MTB/TAN*-derived primary tumor cells were cultured on glass coverslips (Bellco Glass Inc., Cat. # 1943-010015A) and treated with 5µM EdU for 2h before harvest. Coverslips containing cells were fixed in 4% paraformaldehyde for 15 min at room temperature. Cells were then permeabilized in PBS containing 0.5% Triton X-100 for 20 min at room temperature followed by wash steps in 3% bovine serum albumin in 1x PBS before proceeding with the immunofluorescence protocol.

For in vivo immunofluorescence/labeling studies, inguinal #4 mammary fat pads containing primary tumors, residual lesions, or recurrent tumors were dissected and fixed overnight in 4% paraformaldehyde overnight at 4°C. Samples were thoroughly washed and dehydrated prior to paraffin embedding and sectioning at 5µm. Slides were prepared by serial deparaffinization and rehydration followed by PBS washes and antigen retrieval in R-Buffer A (Electron Microscopy Sciences, Cat. # 62706-10) or R-Buffer B (Electron Microscopy Sciences, Cat. # 62706-11) using a retriever (Aptum, Cat. # RR2100-EU).

For GAG immunofluorescence, specificity controls were included where tissue sections were treated with Chondroitinase ABC (Amsbio, Cat. # AMS.E1028-02; 10mU/µl at pH 8) or Heparin lyase (Amsbio, Cat. # AMS.HEP-ENZ III-S; 20mU/µl at pH 7) at 37°C for 1h prior to immunofluorescence.

For labeling studies using coverslips for EdU (Thermo Fisher Scientific, Cat. # C10640) or on tissues for EdU (Sigma Aldrich, Cat. # BCK647-IV-IM-S) or TUNEL (Thermo Fisher Scientific, Cat. # C10619) samples were processed as per manufacturer’s instructions prior to immunofluorescence.

Immunofluorescence samples were blocked with 5% goat serum in 1x PBS with mouse-on-mouse block (Vector Laboratories, Cat. # BMK-2202), washed 3 times in 1x PBS and incubated overnight at 4°C with primary antibodies or matched isotype controls diluted in 5% goat serum in 1x PBS with mouse-on-mouse diluent (Vector Laboratories, Cat. # BMK-2202) as follows: Primary <u>antibodies</u> - Rat anti-Ki67 (Thermo Fisher Scientific, Cat. # 14-5698; 1:100), Rabbit anti-Her2 (Cell Signaling, Cat. # 2165; 1:100), Rabbit anti-cleaved Caspase-3 (Cell Signaling, Cat. # 9664; 1:250), Mouse anti-Chondroitin sulfate (Sigma, Cat. # SAB4200696), Mouse anti-Heparan sulfate (Amsbio, Cat. # 370225), Rabbit anti-GFP (Cell Signaling, Cat. # 2956; 1:200), and Mouse anti-GFP (Living Colors, Cat. # 632381; 1:250). <u>Isotype controls</u> - Rat IgG (Thermo Fisher Scientific, Cat. # 02-9602), Rabbit IgG (Thermo Fisher Scientific, Cat. # 02-6102), Mouse IgG2a kappa (Thermo Fisher Scientific, Cat. # 14-4724-82).

After performing 3 washes with 1x PBS, samples were incubated with secondary antibodies at 37°C for 1h as follows: Secondary antibodies - Goat anti-mouse IgG2a Alexa488 (Thermo Fisher Scientific, Cat. # A21131; 1:1000), Goat anti-rabbit IgG Alexa488 (Thermo Fisher Scientific, Cat. # A11034; 1:1000), Goat anti-rabbit IgG Alexa594 (Thermo Fisher Scientific, Cat. # A11012; 1:1000), Goat anti-rat IgG Alexa568 (Thermo Fisher Scientific, Cat. # A11077; 1:1000), Goat anti-mouse IgM Alexa568 (Abcam, Cat. # ab175702; 1:1000).

Following incubation with secondary antibodies, samples were washed 3 times with 1x PBS and incubated with 0.5µg/ml Hoechst 33258 for 10 min at room temperature for nuclear counter staining. Samples were mounted using ProLong gold (Thermo Fisher Scientific, Cat. # P36934) and imaged using a DM 5000B Automated Upright Microscope with a DFC350 FX monochrome digital camera (Leica Microsystems). Images were quantified using QuPath-0.3.0 open source software.

### Droplet digital PCR

Microdissected lesions from the in vivo experiments or cells harvested from the in vitro dormancy experiment were processed for genomic DNA isolation using the Quick-DNA Midiprep Plus kit (Zymo Research, Cat. # D4075) kit, or the QIAamp DNA mini kit (Thermo Fisher, Cat. # 51304), respectively. To quantify the numbers of GFP^+^ and mCherry^+^ cells, 50ng of gnomic DNA/sample was added together with the ddPCR Supermix for Probes (Bio-Rad, Cat. # 1863025) and the probes of interest: GFP (Bio-Rad, Cat. # dCNS372378948), mCherry (Bio-Rad, Cat. # dCNS507694046), and the control ApoB (Bio-Rad, Cat. # dMmuCNS4075944696). Droplets were generated using the AutoDG system (Bio-Rad, Cat. # 1864101) followed by PCR under the following conditions: 95°C for 5 min; 40 cycles at 94°C for 30 sec and 60°C for 1 min; 98°C for 10 min. Droplets were analyzed using the QX200 Droplet Reader (Bio-Rad, Cat. # 1864003). Droplet-derived copy numbers were first normalized to ApoB numbers to normalize input and then converted to cell numbers by using the average copy number derived from singly transduced cells.

### Western blot

Western blots were performed as described using the following antibodies: β-Tubulin (BioGenex, Cat. # MU122). Secondary antibodies used were anti-mouse 680LT (LI-COR Biosciences 925-68020), anti-rabbit 800CW (LI-COR Biosciences, Cat. # 925-32211),and anti-rat 800CW (LI-COR Biosciences, Cat. # 925-32219). Fluorescent signals were detected using the Odyssey detection system (LI-COR Biosciences), and band intensities were quantified using the Image Studio Ver 2.0 software (LI-COR Biosciences).

### Glycan reductive isotope labeling – mass spectrometry (GRIL-LC/MS)

*MTB/TAN*-derived primary tumor cells were cultured for an in vitro dormancy time course as previously described and harvested at baseline (D0), deinduction (D7, D28), and reinduction (D28+) time points. At the time of harvest, cells were kept on ice, thoroughly washed 2 times with ice cold 1x PBS, and gently scraped into 1x PBS. Cells were pelleted down at 1500 rpm and the pellets were flash frozen for further analysis.

GAG isolation and purification was performed by sonicating the cell pellet in ultrapure distilled water (Invitrogen, Cat. # 10977-015) containing measured amount of protease inhibitor cocktail SetIII, EDTA free (EMD Millipore, Cat. # 539134-1ml) followed by adding equal volume of 2X wash buffer (100mM NaOAc and 400mM NaCl; pH 6). Protein digestion was done using bacterial protease (Sigma, Cat. # P5147) at a concentration of 0.4mg/ml at 37°C overnight (16h). Samples were then loaded on to a Poly-Prep column (Bio-Rad, Cat. # 731-1550) packed with pre-equilibrated DEAE Sephacel gel (Sigma, Cat. # 16505). Columns were washed with 10 bed volume of wash buffer (50mM NaOAc containing 200mM NaCl; pH 6) and bound GAG were eluted using elution buffer (50mM NaOAc containing 1M NaCl; pH 6). Eluted samples were then loaded on to a PD10 desalting column (GE Healthcare, Cat. # 17-0851-01) pre-washed with 10% ethanol. Desalted GAG samples in 10% ethanol were lyophilized and used for further analysis.

Lyophilized samples were resuspended in MilliQ water and distributed for enzymatic digestion of chondroitin sulfate and heparan sulfate at 37°C for 24h using 30mU of Chondroitinase ABC (Amsbio, Cat. # AMS.E1028-02) and 10mU of Heparinase I, II, III (Ibex, Cat. # PN 50-010; PN 50-011; PN 50-012 Heparinase I, II, and III), respectively. Digested samples were spin-filtered using a 3K MWCO spin filtering unit (Pall Life Sciences, Cat. # OD003C34) and tagged with ^12^C_6_-aniline in the presence of a reducing agent 1M NaCNBr in a mixture of DMSO: HOAc (65:35 v/v). Reductive isotope labeling is performed at 65°C for 45 minutes, followed by incubation at 37°C for 16 h. Samples were mixed with internal standards (^13^C_6_-aniline tagged disaccharides from chondroitin sulfate and heparan sulfate respectively) and LC/MS is performed in negative mode using an LTQ-Orbitrap (Thermo Fisher Scientific) mass spectrometer.

### Statistics

For each gene expression signature in each data set, signature scores were calculated as weighted averages of z-score transformed expression data across signature genes, where the z-score transformation was done for each gene across the samples, and the weights are 1 for genes expected to be positively associated with the signature target (e.g. dormancy) and -1 for genes expected to be negatively associated with the signature target.

CRISPR screen sequencing reads were de-convoluted using the unique barcodes associated with each sample and mapped to sgRNA sequences. The abundance of each sgRNA was normalized by the median of ratios (MoR, as implemented in DESeq2), the total read count (T, as % of total), the robust z-score (RZ, with median-centering and MAD-scaling), trimmed mean of M-values (TMM, as implemented in edgeR), and normalized rank (NR, as percentile rank) following which depletion and enrichment scores for each sgRNA were calculated by MAGeCK or the Mann-Whitney U test followed by robust rank aggregation (RRA). False positive rates were controlled for using data from the size matched non-targeting negative control (NC) sgRNA library. In addition to the five normalization methods above, an NC-centric version of MoR was done using negative control gRNAs to calculate scaling factors instead of using all gRNAs, and an NC-centric version of RZ was done using the median and MAD values calculated from negative control gRNAs instead of all gRNAs. Methods with the highest sensitivity were as follows: MAGeCK-RRA-M, MAGeCK-RRA-T, MW-RRA-MoR, MW-RRA-NC_MoR, MW-RRA-NC_RZ, MW-RRA-NR, MW-RRA-RZ, MW-RRA-TMM. Using *p* values from the 8 tests above, hits called by a larger number of tests were prioritized for further validation.

To determine differences between multiple groups, ANOVA followed by Tukey’s multiple comparisons test was used. In cases where the data were not normally distributed, the Mann-Whitney U test was used to determine statistical significance. Survival curves were subjected to Kaplan-Meier analysis and *p* values and hazard ratios were calculated using the Mantel-Haenszel method.

---

**Fig. S1:**
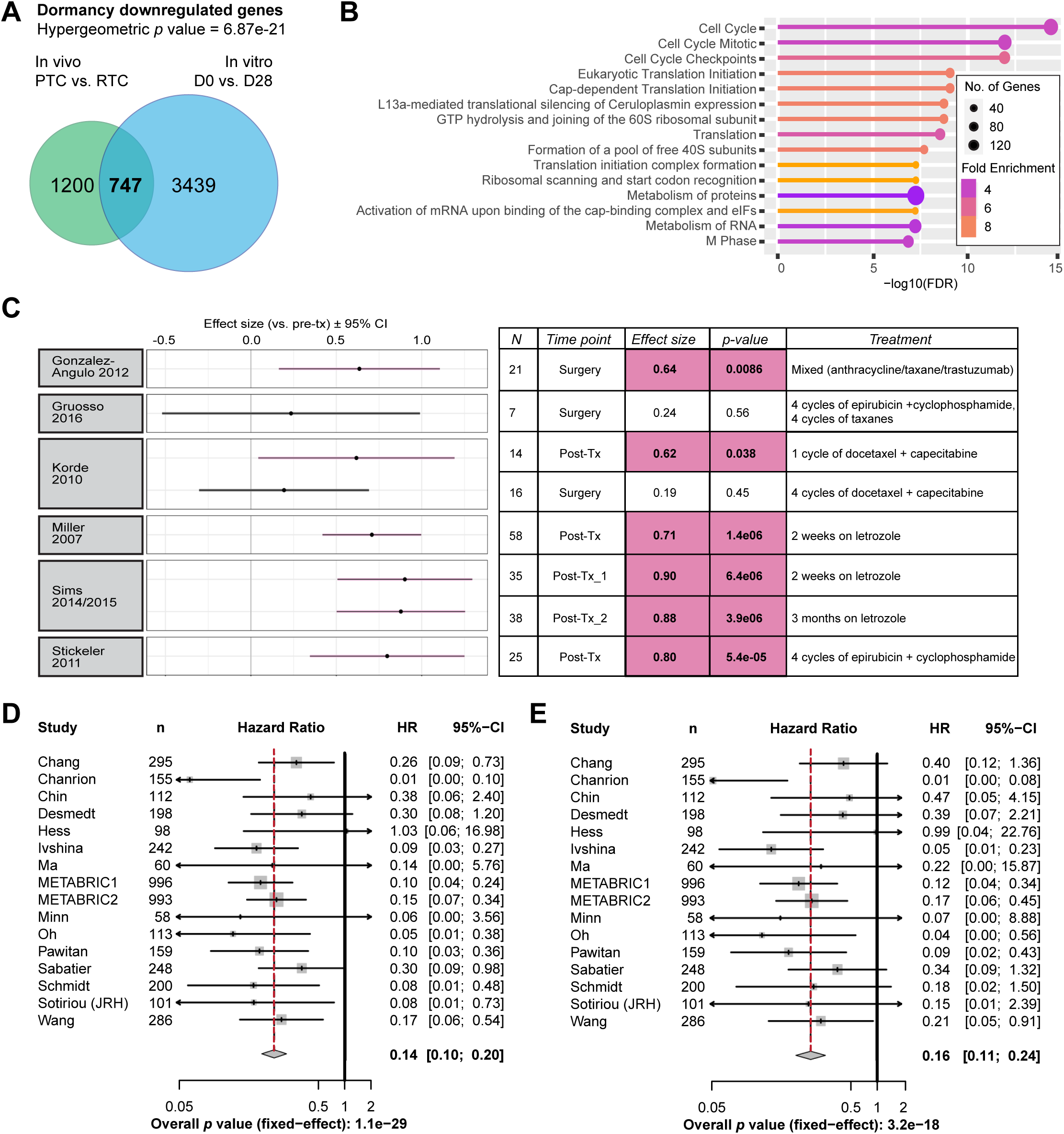
**A.** Hypergeometric test and Venn diagram depicting the overlap between the in vivo and IVD-derived downregulated gene set. **B.** Top 15 Reactome gene ontology terms for the overlapping upregulated set of 747 genes is depicted on the bar plot. **C.** Dormancy-associated tumor cell-autonomous gene signature (after subtracting out genes that belong to the gene ontology term ‘Cell Cycle’) enrichment following neoadjuvant therapies vs. pre-treatment (pre-tx) samples across 7 different datasets is measured by effect size. Significant effects are highlighted in pink. **D.** Forest plot representation of hazard ratios (HR) and 95% confidence intervals (CI) as a function of the dormancy-signature or **E.** dormancy signature after subtracting out genes that belong to the gene ontology term ‘Cell Cycle’ in breast cancer patients recurring 10 years after initial treatment. Red dashed lines depict the shift in HR across 16 human datasets. Forest plot representation of hazard ratios (HR) and 95% confidence intervals (CI) as a function of the dormancy-signature in breast cancer patients recurring 10 years after initial treatment.

**Fig. S2:**
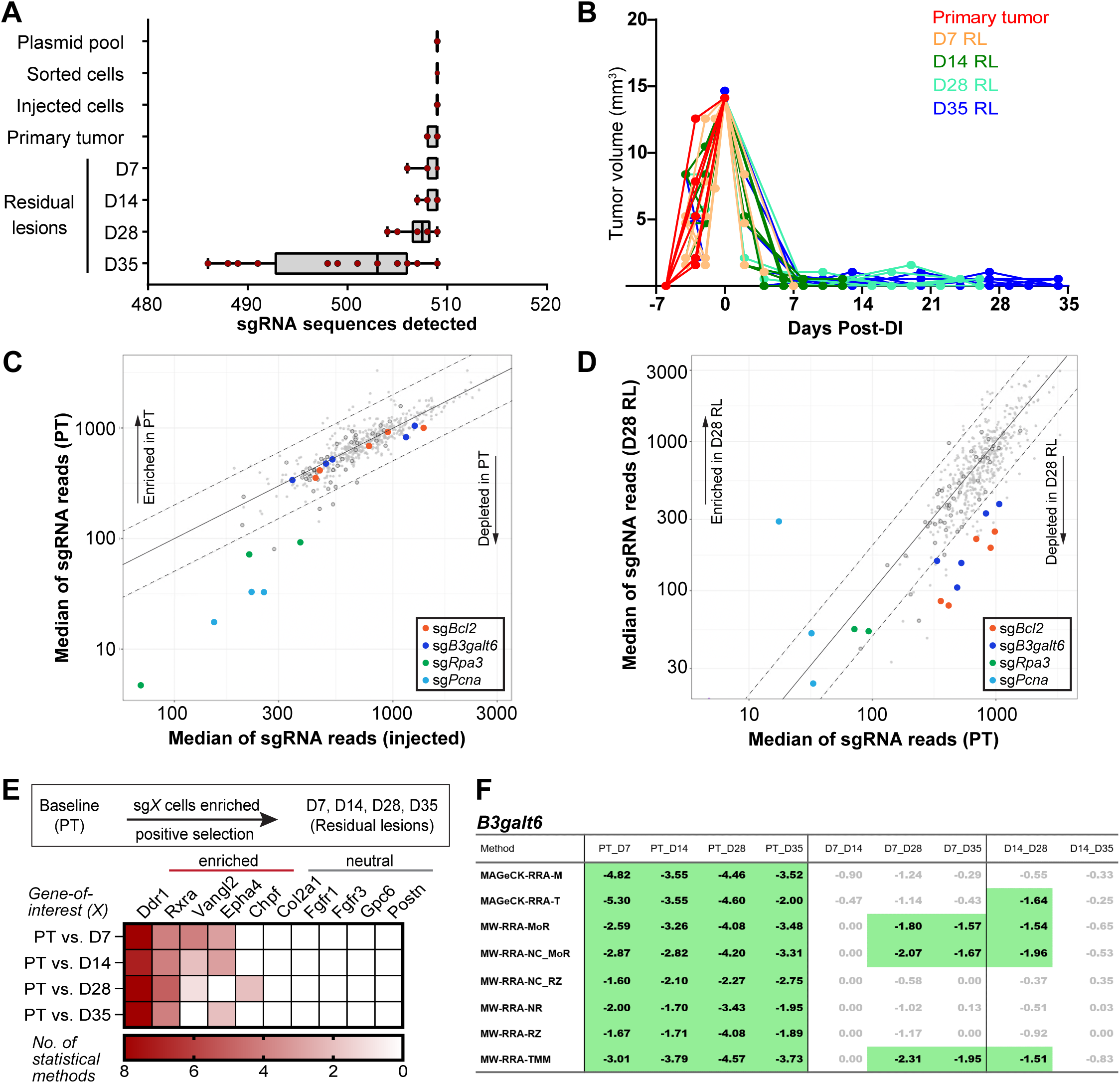
**A.** Number of sgRNAs (out of 509) detected in every sample at sequential time points in the CRISPR-Cas9 screen. Data are represented as median ± interquartile range. **B.** Palpation data tracking tumor volumes for all harvested samples at PT and RL time points. **C, D** Scatterplot of sgRNA sequence reads normalized by MoR in injected vs. PT samples (**C**) and PT vs. D28 RL (**D**). Solid line indicates identity line, dotted lines indicate two-fold differences. Outlined grey circles indicate non-targeting sgRNAs. **E.** Subset of CRISPR-Cas9 screen enrichment hits (5 genes) identified by 2-8 statistical methods (MAGeCK-RRA-M, MAGeCK-RRA-T, MW-RRA-MoR, MW-RRA-NC_MoR, MW-RRA-NC_RZ, MW-RRA-NR, MW-RRA-RZ, MW-RRA-TMM) and neutral genes (5 genes) displaying no selection during dormancy. **F.** Scores indicate –log_10_(p value) for each pairwise comparison using PT, D7, or D14 as a baseline. Rows represent the statistical method being used. Positive values indicate enrichment of sgRNAs, negative values indicate depletion of sgRNAs. Significant results are highlighted in green.

**Fig. S3:**
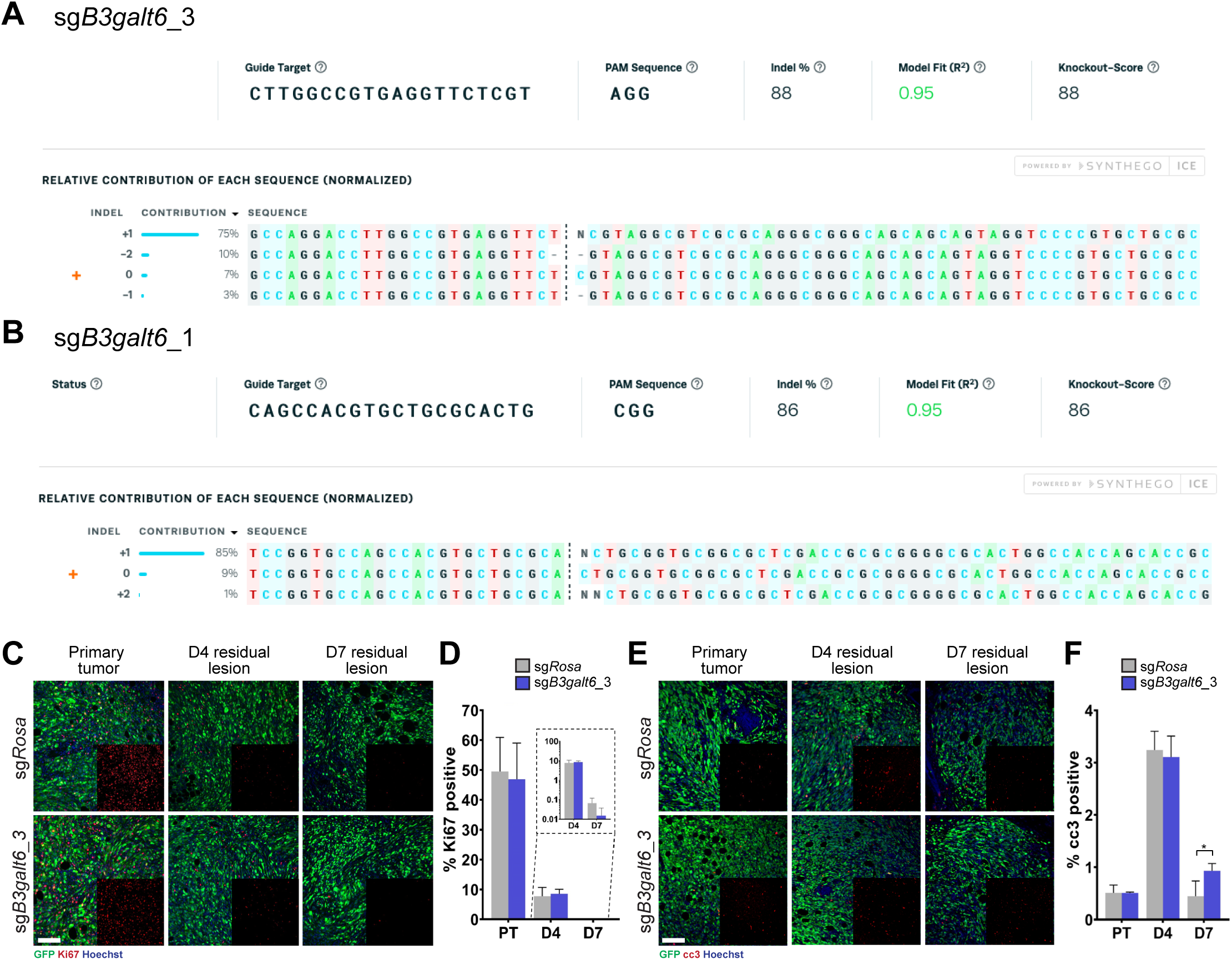
**A.** Inference of CRISPR edits (ICE) analysis for sg*B3galt6*_3 and **B.** sg*B3galt6*_1 displaying the proportion of indels in the population. Wild-type sequence is represented by the orange + sign at 0. Dotted line indicates the Cas9 cut site. **C.** Immunofluorescence and **D.** associated quantification for Ki67 (red) or **E, F.** cleaved caspase 3 (cc3, red) in Her2-dependent-Cas9 sg*Rosa* and sg*B3galt6*_3 (green) PTs, D4, and D7 RLs. Scale bar=100μm. Quantification in the sg*Rosa* (grey) and sg*B3galt6*_3 (dark blue) groups is represented as mean ± SD. \**p*<0.05.

**Fig. S4:**
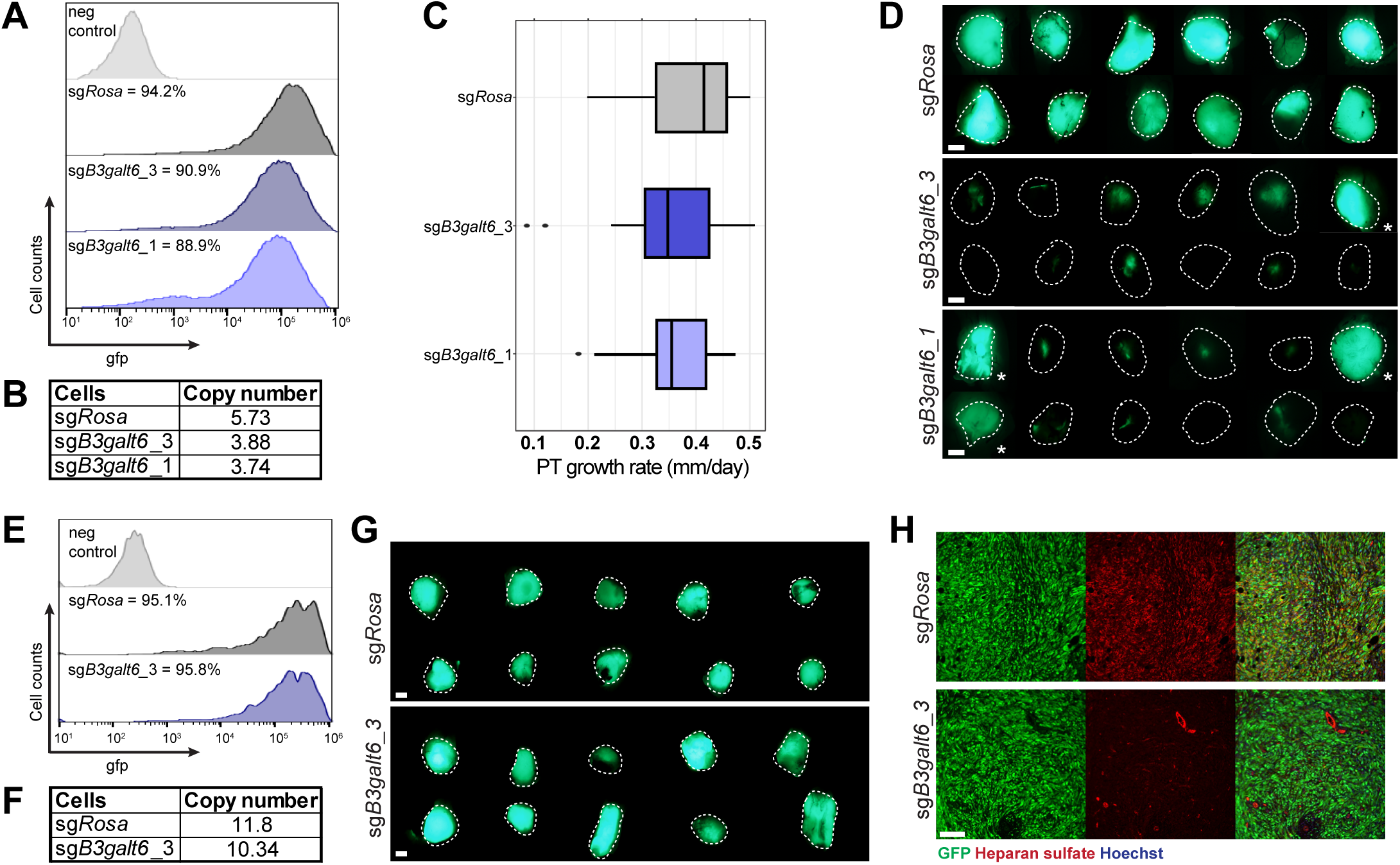
**A.** Histograms represent the distribution of GFP positivity in the injected Her2-dependent-Cas9 sg*Rosa* (dark grey), sg*B3galt6*_3 (dark blue), and sg*B3galt6*_1 (light blue) relative to untransduced Her2-dependent-Cas9 cells (light grey). **B.** Average copy number of integrated GFP in the injected Her2-dependent-Cas9 sg*Rosa*, sg*B3galt6*_3, and sg*B3galt6*_1 cells detected by ddPCR. **C.** Growth rate of primary sg*Rosa* (dark grey), sg*B3galt6*_3 (dark blue), and sg*B3galt6*_1 (light blue) tumors. Data are represented as median ± interquartile range. **D.** Stereoscope images of 12 representative recurrences harvested. Dotted white lines drawn based on discernible tumor edges in bright field images. White asterisks indicate GFP+ recurrences in the sg*B3galt6*_3 and sg*B3galt6*_1 groups. Scale bar=2mm. **E.** Histograms represent the distribution of GFP positivity in the injected Her2-dependent-Cas9 sg*Rosa* (dark grey) and sg*B3galt6*_3 (dark blue) relative to untransduced Her2-dependent-Cas9 cells (light grey). **F.** Average copy number of integrated GFP in the injected Her2-dependent-Cas9 sg*Rosa* and sg*B3galt6*_3 cells detected by ddPCR. **G.** Stereoscope images of 10 representative recurrences harvested. Dotted white lines drawn based on discernible tumor edges in bright field images. Scale bar=2mm. **H.** Immunofluorescence for heparan sulfate (red) in Her2-dependent-Cas9 sg*Rosa* or sg*B3galt6*_3 (green) recurrences. Scale bar=100µm.

**Fig. S5:**
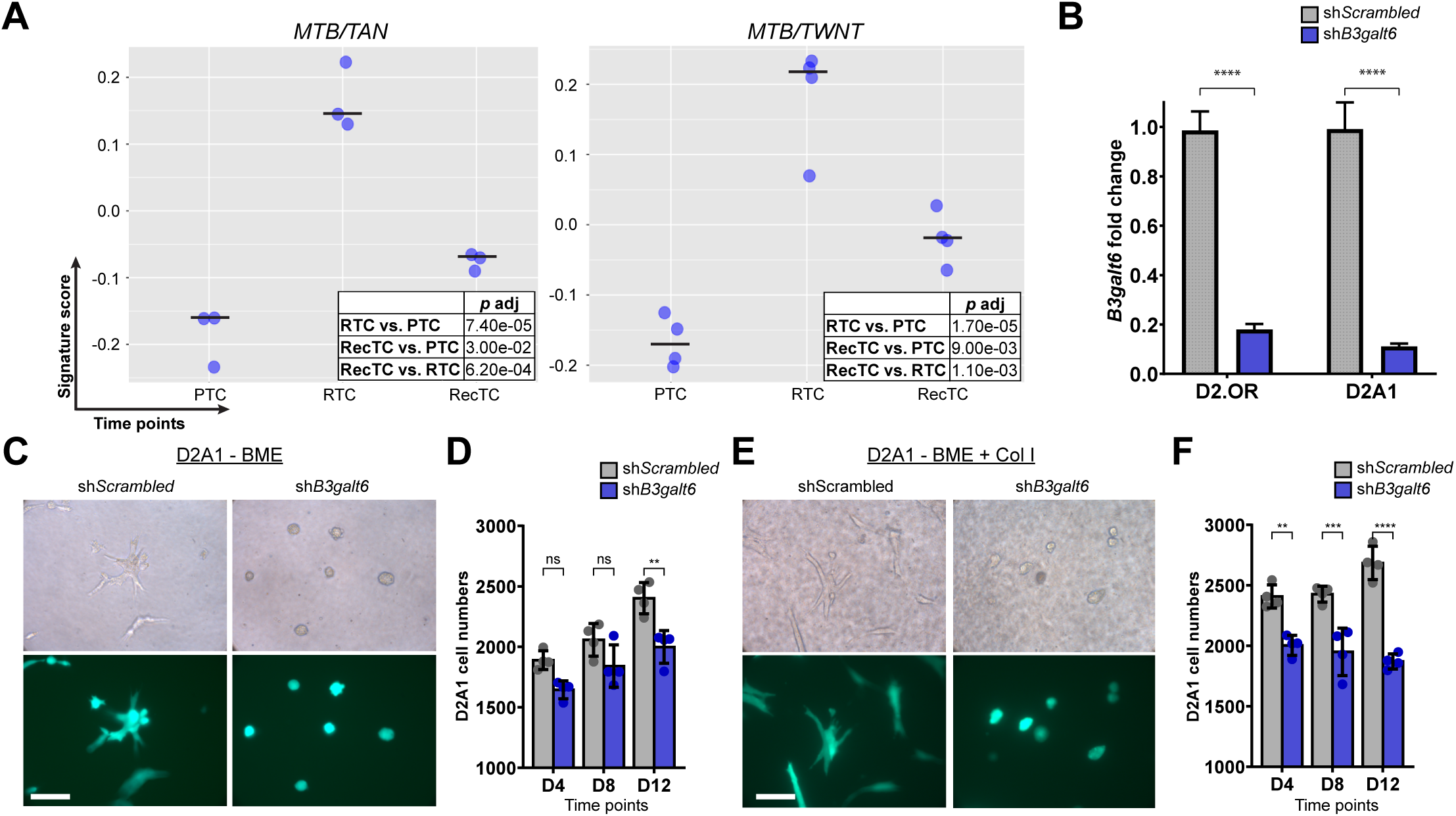
**A.** Application of the gene expression signature derived from D2.OR (dormant, indolent) cells vs. D2A1 (proliferative, aggressive) cells in 3D to the *MTB/TAN* or *MTB/TWNT*-derived in vivo dormancy temporal profiling. Table represents adjusted *p* value for each pairwise comparison between primary tumor cells (PTC), residual tumor cells (RTC), and recurrent tumor cells (RecTC). Black lines indicate median score for each time point. **B.** qRT-PCR for *B3galt6* transcripts in sh*Scrambled* or sh*B3galt6* D2.OR and D2A1 cells. **C.** Brightfield and fluorescence images of sh*Scrambled* and sh*B3galt6* D2A1 cells grown in 3D on basement membrane extract (BME) and **D.** associated viable cell numbers measured by absorbance at 570nm at D0, D8, and D12 time points. Scale bar=100µm. Data are represented as mean ± SD. ns=non-significant, \*\**p*<0.01. **E.** Brightfield and fluorescence images of sh*Scrambled* and sh*B3galt6* D2A1 cells grown in 3D on BME + Col I and **F.** associated viable cell numbers measured by absorbance at 570nm at D0, D8, and D12 time points. Scale bar=100µm. Data are represented as mean ± SD. \*\**p*<0.01, \*\*\**p*<0.001, \*\*\*\**p*<0.0001.

**Fig. S6:**
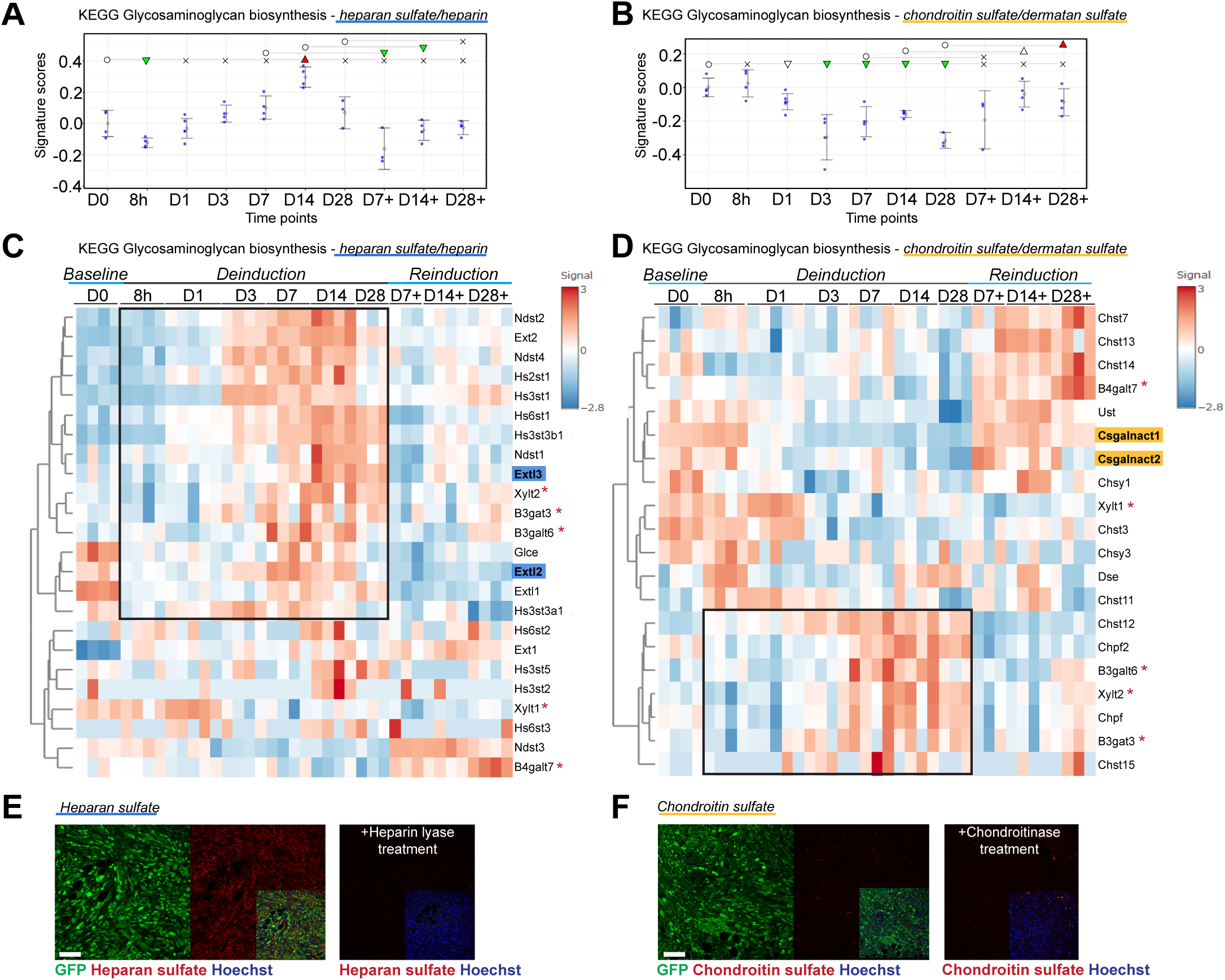
**A.** Application of KEGG glycosaminoglycan (GAG) biosynthesis signatures for heparan sulfate/heparin and **B.** chondroitin sulfate/dermatan sulfate on MTB/TAN IVD temporal gene expression profiles. Circle=baseline, x=non-significant, empty triangles=trending significance 0.05<*p*<0.1, filled triangles=*p*<0.05, inverted triangles=decreased signature score, upright triangles=increased signature score. **C.** Heatmap of KEGG glycosaminoglycan (GAG) biosynthesis gene expression signatures for heparan sulfate/heparin (left) and **D.** chondroitin sulfate/dermatan sulfate (right) on *MTB/TAN* IVD temporal gene expression profiles. Black box represents the major group demonstrating dormancy-selective upregulation identified by hierarchical clustering. Red asterisks represent enzymes involved in tetrasaccharide linker synthesis during proteoglycan assembly. Blue boxes indicate enzymes involved in determining heparan sulfate synthesis, yellow boxes indicate enzymes involved in chondroitin sulfate synthesis. **E.** Immunofluorescence for heparan sulfate (left panels) or **F.** chondroitin sulfate (right panels) (red) in PTs derived from Her2-dependent-Cas9 cells with sg*Rosa* (green). Heparan sulfate/Chondroitin sulfate staining following heparin lyase or chondroitinase treatment is used as the negative control. Scale bar=100µm.

**Fig. S7:**
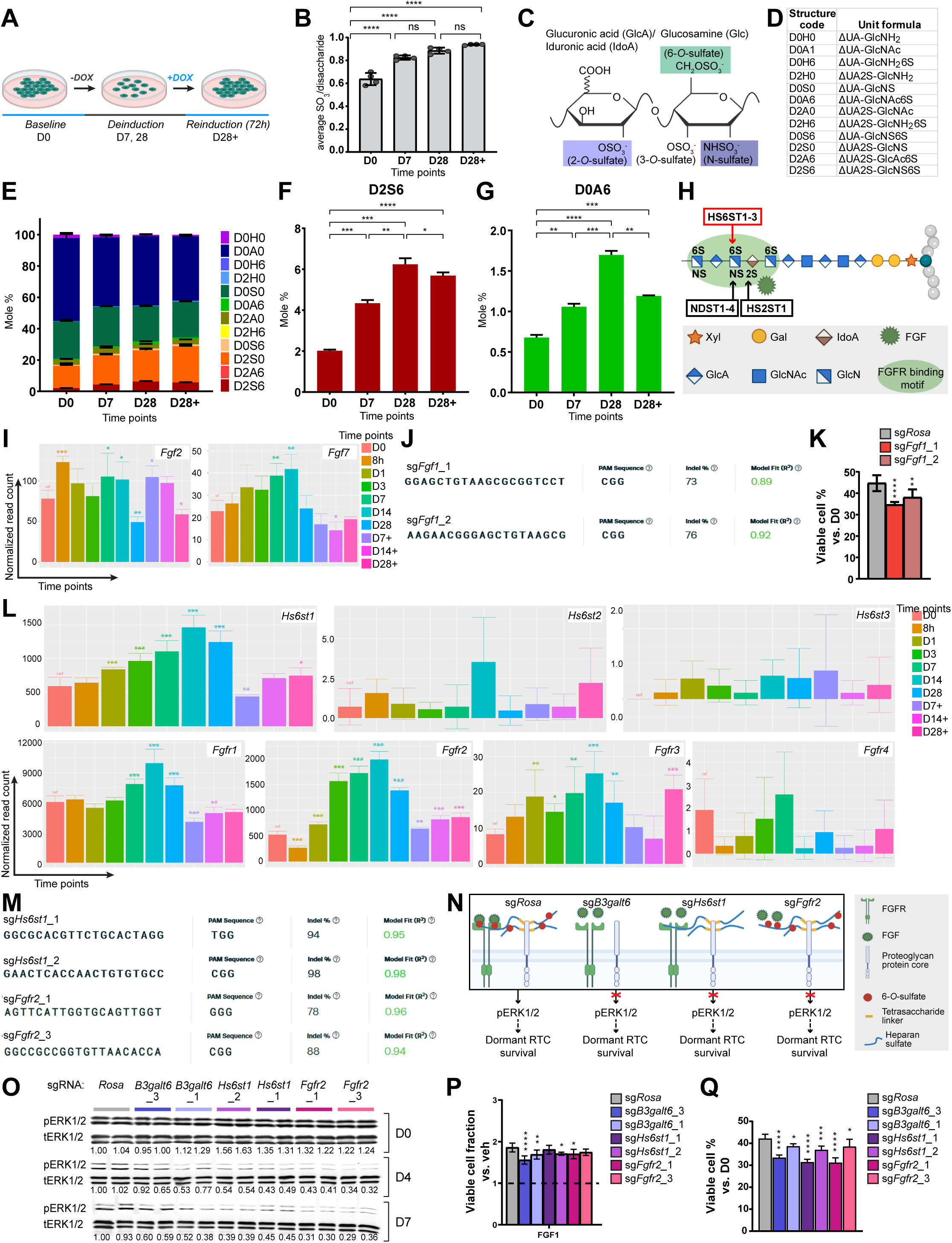
**A.** Experimental setup for IVD samples isolated for LC/MS for heparan sulfate and chondroitin sulfate disaccharide analysis. **B.** Average sulfation/disaccharide of heparan sulfate at D0 (*baseline*), D7, D28 (*deinduction*), and D28+ (*reinduction*) time points measured by LC/MS. Data are represented as mean ± SD. ns=non-significant, \*\*\*\**p*<0.0001. **C.** Possible sites of sulfation on the heparan sulfate disaccharide. Colored boxes indicate the types of sulfation assessed. **D.** 4-character disaccharide structure code (DSC) nomenclature and unit formulae. ΔUA – Δ4,5-unsaturated uronic acid. **E.** Distribution of heparan sulfate disaccharide motifs at IVD time points represented according to the 4-character disaccharide structure code (DSC) nomenclature. Molar percentages of (**F**) D2S6 and (**G**) D0A6 are highlighted. Data are represented as mean ± SD. ns=non-significant, \**p*<0.05, \*\**p*<0.01, \*\*\**p*<0.001, \*\*\*\**p*<0.0001. **H.** Pictorial representation of the substrates and the type of sulfation catalyzed by sulfotransferase enzymes to generate protein binding motifs, *e.g.*, FGFR binding motif. **I.** Normalized read counts indicating the expression of *Fgf2* and *Fgf7*. Asterisks indicate significant changes in normalized read counts vs. D0 (*baseline*) \**p*<0.05, ** *p*<0.01. **J.** Inference of CRISPR edits (ICE) analysis for sg*Fgf1*_1 and sg*Fgf1*_2 displaying the proportion of indels in the population. **K.** Viable RTC counts relative to D0 in sg*Rosa*, sg*Fgf1*_1, and sg*Fgf1*_2 Her2-dependent-Cas9 cells. \*\**p*<0.01, \*\*\*\**p*<0.0001. **L.** Normalized read counts indicating the expression of *Hs6st1-3* and *Fgfr1-4*. Asterisks indicate significant changes in normalized read counts vs. to D0 (*baseline*) \**p*<0.05, \*\**p*<0.01, \*\*\**p*<0.001. **M.** Inference of CRISPR edits (ICE) analysis for sg*Hs6st1*_1, sg*Hs6st1*_2, sg*Fgfr2*_1, sg*Fgfr2*_3 displaying the proportion of indels in the population. **N.** Schematic depicting the role of heparan sulfate, 6-*O*-sulfation, and FGF signaling in maintain dormant RTC survival. **O.** Western blot quantification of phospho-ERK1/2 (pERK1/2) over total ERK1/2 (tERK1/2) levels in sg*Rosa*, sg*B3galt6*_3, sg*B3galt6*_1, sg*Hs6st1*_2, sg*Hs6st1*_1, sg*Fgfr2*_2, and sg*Fgfr2*_3 Her2-dependent-Cas9 cells at D0 (*baseline*), D4, and D7 (*deinduction*) time points. pERK1/2/tERK1/2 signal normalized to sg*Rosa* levels at D0. Asterisks indicate significant changes vs. sg*Rosa* signal within the time point (*baseline*) \**p*<0.05, \*\**p*<0.01, \*\*\**p*<0.001. **P.** Viable RTC counts relative to vehicle-treated cells (black dashed line) at D4 in sg*Rosa*, sg*B3galt6*_3, sg*B3galt6*_1, sg*Hs6st1*_2, sg*Hs6st1*_1, sg*Fgfr2*_1, and sg*Fgfr2*_3 Her2-dependent-Cas9 cells treated with FGF1 (25ng/ml). Black dotted line indicates the viable cell number in the vehicle controls. \**p*<0.05, \*\**p*<0.01, \*\*\*\**p*<0.0001. **Q.** Viable RTC counts relative to D0 in sg*Rosa*, sg*B3galt6*_3, sg*B3galt6*_1, sg*Hs6st1*_2, sg*Hs6st1*_1, sg*Fgfr2*_1, and sg*Fgfr2*_3 Her2-dependent-Cas9 cells. \**p*<0.05, \*\*\*\**p*<0.0001.

